# Analysis of bacterial genomes from an evolution experiment with horizontal gene transfer shows that recombination can sometimes overwhelm selection

**DOI:** 10.1101/186759

**Authors:** Rohan Maddamsetti, Richard E. Lenski

**Affiliations:** Ecology, Evolutionary Biology, and Behavior Program, Michigan State University, East Lansing, MI; BEACON Center for the Study of Evolution in Action, Michigan State University, East Lansing, MI; Department of Systems Biology, Harvard Medical School, Boston, MA; Department of Microbiology and Molecular Genetics, Michigan State University, East Lansing, MI

## Abstract

We analyzed genomes from an experiment in which *Escherichia coli* K-12 Hfr donors were periodically introduced into 12 evolving populations of *E. coli* B. Previous work showed that recombination did not increase adaptation, despite increasing variation relative to asexual controls. The effects of recombination were highly variable: one lineage was mostly derived from the donors, while another acquired almost no donor DNA. In most lineages, some regions showed repeated introgression and others almost none. Regions with high introgression tended to be near the donors origin of transfer sites. To determine whether introgressed alleles imposed a genetic load, we extended the experiment for 200 generations without recombination and sequenced whole-population samples. Beneficial alleles in the recipient populations were occasionally driven extinct by maladaptive donor-derived alleles. On balance, our analyses indicate that the plasmid-mediated recombination was sufficiently frequent to drive donor alleles to fixation without providing much, if any, selective advantage.

## Introduction

An open question in microbial evolution is why some populations of bacteria seem to have extensive intergenomic recombination (Rosen *et al.* 2015) while others seem to have little (Sarkar and Guttman 2004). Some symbiotic bacteria appear to be entirely clonal owing to their tight associations with hosts that preclude contact with other lineages (Moran *et al.* 2009), but otherwise the reasons for differences in recombination rates across bacterial taxa are unclear. Intergenomic recombination can potentially break the linkage between particular beneficial or deleterious mutations and the rest of the genome. Under conditions of high recombination and strong selection, individual genes, rather than entire genomes, can thus go to fixation. When recombination is infrequent or absent but selection is strong, highly beneficial mutations can drive large genomic regions or even whole genomes to fixation. Recent work shows that both gene-specific and genome-wide selective sweeps occur in microbial communities (Bendall *et al.* 2016). In contrast to meiotic recombination, however, bacterial recombination replaces a recipient allele with a donor allele, rather than swapping homologous regions between chromosomes. Thus, recombination in bacteria does not necessarily break up linkage disequilibrium across distant sites on the recipient chromosome; instead, it may preserve a “clonal frame” over most of the chromosome, interrupted by stretches of DNA introduced by horizontal gene transfer (Milkman and Bridges 1990).

Horizontally transmitted viruses and conjugative plasmids mediate recombination in many species of bacteria (Levin and Lenski 1983). In this context, recombination is a special kind of evolutionary process. Like mutation rates, intergenomic recombination rates can evolve because many gene products involved in DNA replication and repair affect the rates of mutation and recombination. But unlike mutation rates, recombination rates in bacteria are also subject to coevolution owing to the association with plasmids and viruses. Intergenomic recombination qualitatively changes evolutionary dynamics and can speed up adaptive evolution under some circumstances (Keightley and Otto 2006; Cooper 2007; McDonald *et al.* 2016). However, recombination—especially as it occurs in bacteria—may have originated as a byproduct of the spread of selfish elements rather than as a means of increasing the efficiency of natural selection in the genome as a whole (Hickey and Rose 1986).

In this paper, we revisit an evolution experiment in which recombination did not appear to increase the efficiency of natural selection, nor did it speed up adaptation. Souza, Turner, and Lenski (Souza *et al.* 1997) conducted an experiment in which they periodically introduced strains of *Escherichia coli* K-12 that could donate genetic material but not grow (owing to mutations that caused nutritional deficiencies) into populations of *E. coli* B that had previously evolved in and adapted to a glucose-limited minimal medium for 7000 generations. We refer to the recombination treatment in that study as the Souza-Turner-Lenski experiment (STLE) and to the experiment that generated the asexual recipients as the long-term evolution experiment (LTEE). The goal of the STLE was to test whether recombination would increase the rate of adaptation by increasing the genetic variation available to natural selection (Souza *et al.* 1997). In the absence of complex selection dynamics (such as frequency-dependent selection), the expected rate of adaptation of a population is proportional to the genetic variance in fitness in a population (Fisher 1930). The outcome of the STLE was unexpected in that recombination demonstrably introduced substantial genetic variation (as determined by tracking ~10 genetic markers available to the authors at that time), but it had no significant effect on the rate of adaptation compared to control populations that evolved without recombination. Two possible hypotheses were proposed that might explain those results. According to one, the recombination treatment was so effective that it acted like a strong mutational force, replacing many neutral alleles (or even overwhelming selection, if donor alleles were maladaptive) through the sheer flux of donor DNA into the recipient genomes. Alternatively, the interactions between donor and recipient genes, and the ecological context in which those interactions occurred, might have somehow generated strong frequency-dependent selection, such that the assays used to measure fitness gains were undermined. In fact, one recombinant population of the STLE actually appeared to decline substantially in fitness, an effect that was shown to reflect an evolved frequency-dependent interaction (Turner *et al.* 1996). In this study, we characterize the genomic evolution that occurred during the STLE, with the aim of resolving the puzzling outcome of the STLE. On balance, we find evidence that the recombination treatment acted more like a greatly increased mutational force than like a sieve separating beneficial from deleterious mutations.

## Methods

### Overview of the STLE

The experiments performed by Souza *et al.* (1997) are described fully in that paper. In brief, 12 recombinant populations and 12 control populations were started from clones isolated after 7000 generations of the LTEE (Table 1). These populations were propagated daily for 1000 generations (150 days) following the same transfer regime and using the same DM25 medium and temperature as the LTEE (Lenski *et al.* 1991). However, on day 0 and every fifth day thereafter (~33 generations), a mixture of four K-12 Hfr donor strains (REL288, REL291, REL296, and REL298) was added to the recombination treatment populations and allowed to conjugate for 1 h. The ratio of donor to recipient cells during the conjugation treatment was ~4:1. All four donors were auxotrophic for one or more essential amino acids (arginine, leucine, or isoleucine-valine), so they could transfer their genetic material but not grow and persist in the population.

**Table 1.**
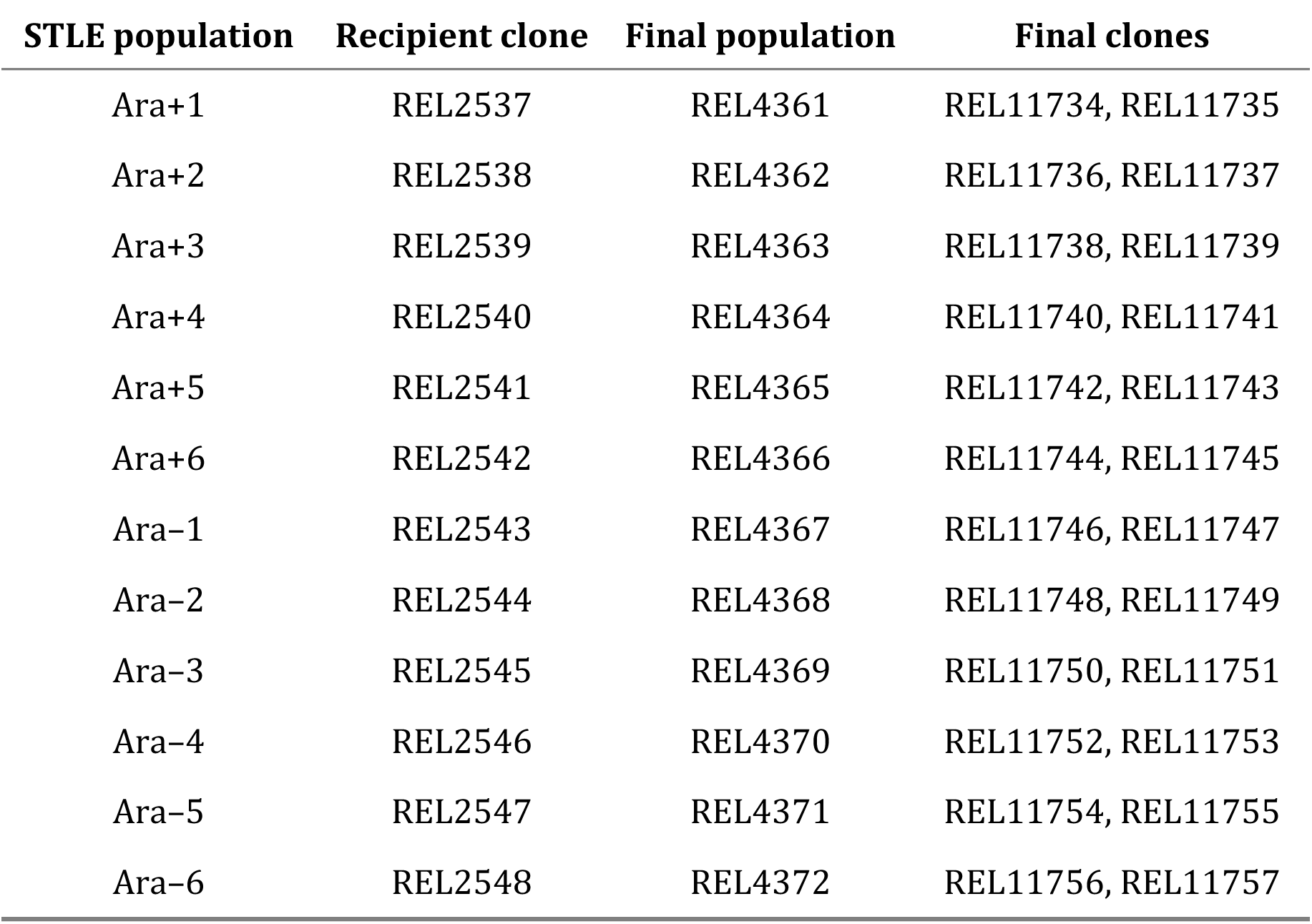
The identifying numbers and relationships of the 12 sequenced recipient clones used to start the 12 STLE populations and the 24 recombinant clones isolated at the end of the STLE.

### Isolation of clones for genomic analysis

Samples of all 12 recombination-treatment populations from generation 1000 of the STLE (Table 1) were revived as follows: 100 μL from each frozen stock were pipetted into 10 mL of LB medium, grown for 24 h, diluted and grown in DM25 for two more 24 h cycles, and then spread on LB agar plates. The two colonies that grew closest to randomly placed marks were re-streaked on LB agar plates and then grown in LB medium. These 24 STLE-derived recombinant clones were stored at -80°C (Table 1). We sometimes refer to these clones as “odd” and “even” (based on the freezer identification numbers for each pair) when presenting data for just one clone from each population.

### Genome sequencing and analysis

The 4 Hfr donor strains (REL288, REL291, REL296, and REL298), 12 LTEE-derived recipient clones used as ancestors in the STLE, and 24 STLE-derived recombinant clones (two per population) were thawed and grown in LB medium, and samples of genomic DNA were isolated from each one. The genomic DNA was then sequenced on an Illumina MiSeq at the MSU RTSF Genomics Core Facility. We used the *breseq* (version 0.31) pipeline (Deatherage *et al.* 2015) to analyze the genomes, with the ancestral LTEE strain REL606 as the reference genome unless otherwise specified. We used the *gdtools* utility in *breseq* to compute a table that lists the union of all the mutations found in the K-12 donor genomes in comparison to REL606, and to identify mutations specific to each of the donor strains.

A custom python program called *label_mutations.py* was written that performs four tasks. First, it labels all of the mutations (i.e., all genetic differences) found in the STLE-derived recombinant genomes relative to the REL606 genome by looking at the same site in the donor, recipient, and recombinant genomes. Nine distinct labels are used, as follows. (i) Mutations that are present in both a recombinant clone and its parent recipient clone (but not in REL606 or the union of donor strains) are labeled as “LTEE mutations”. (ii) Mutations found in both a recombinant clone and the union of K-12 donors are labeled as “K-12 mutations”; they represent horizontally transferred alleles. (iii) Any mutations that are present in a recombinant clone, but not found in either the donors or recipient clone, are labeled as “new mutations”; these are mutations that occurred during the STLE, and which do not appear to involve a donor. (iv) Mutations present in the union of K-12 donors that are not found in the recipient clone are labeled “REL606 mutations;” more precisely, these sites are genetic markers that distinguish the *E. coli* B-derived REL606 strain used to start the LTEE from K-12. Supplementary Figure S1 shows the distribution of these markers along the *E. coli* B chromosome. (v). Mutations found in a recipient, but not in its derived recombinant, are labeled as “deleted mutations”; these mutations were removed by recombination with the donors or otherwise lost during the STLE. (vi-ix). Mutations specific to each of the four donor strains that are found in the recombinants are labeled as “donor-specific mutations”; these mutations are included in the set of “K-12 mutations” in most of our analyses. The program ignores all sites that are identical between the recombinant, recipient, and donors because they provide no useful information. We also ignore genomic “islands” that are present in K-12 but not REL606. As a second task, the *label_mutations.py* program produces a table of the genetic markers that distinguish K-12 and REL606 (see label iv above). Third, this program generates a table of the LTEE-specific mutations found in the recipient genomes (see labels i and v above).

An R script called *dissertation-analysis*.R makes figures and performs statistical tests using the tables of labeled mutations that *label_mutations.py* produces.

### Manual annotation of specific donor genome features

Each donor strain also has two special features: the mutations that made it an amino-acid auxotroph, and the Hfr transfer origin and orientation (i.e., direction of transfer). The *breseq* analysis (using version 0.31) found all the auxotrophy mutations present in the donor strains REL288, REL291, REL296, and REL298. These mutations were generated and annotated by Wanner *et al.* (1986), and the strains were confirmed to be auxotrophs by Souza *et al.* (1997). We checked the genomic locations of the auxotrophy mutations returned by *breseq* against the original strain annotations and a fine-grained traditional linkage map for *E. coli* K-12 from the Coli Genetic Stock Center (https://cgsc2.biology.yale.edu/Workingmap.php, last accessed 7/13/17; Berlyn 1998). We note that the *leuB* auxotrophy mutation in REL298 is a S286L (TCG→TTG) mutation in an NAD-binding region of the protein (http://www.uniprot.org/uniprot/P30125, last accessed 7/31/17), rather than an amber nonsense mutation as originally annotated.

We did not resolve the position and orientations of the Hfr transfer origins, because the F plasmid contains repetitive sequences and insertion elements that map to multiple locations on the *E. coli* chromosome. The genomes of K-12 and REL606 are largely syntenic (Studier *et al.* 2009), and so we used the K-12 linkage map to place the Hfr *oriT* transfer origin sites in the K-12 donors with respect to their homologous genes in REL606. This mapping of K-12 elements to the REL606 genome is only approximate, but it appears to perform reasonably well. Our annotations using *breseq* and the K-12 linkage map along with cross-references to annotations from the Coli Genetic Stock Center are in a file called *donor_hand_annotations.csv*.

### Calculation of lengths of donor and recipient segments in recombinant genomes

Recombination breakpoints occur somewhere in the interval between donor-specific and recipient-specific markers. A minimum estimate of the length of a donor segment would place the recombination breakpoints at the donor markers on each end, while a maximum estimate would place the breakpoints at the flanking recipient markers. In fact, the true breakpoints cannot be known exactly. Our approach uses the minimal estimate on the left, but the maximal estimate on the right, and so it will produce intermediate values that should tend to an overall average length similar to what would be obtained by averaging the minimum and maximum segment lengths. In particular, our algorithm alternates between K-12 and B-derived segments along the genome, switching states when it reaches the alternate marker type. This algorithm is described in detail in the Supplement.

### STLE continuation experiment

We performed the following experiment to determine the fate of introgressed K-12 alleles in the absence of the recombination treatment. We revived the 12 populations of the STLE recombination treatment from the samples that were frozen at 1000 generations. We then propagated them daily for 200 generations (30 days) following the same transfer regime and using the same DM25 medium and other conditions as the LTEE (Lenski *et al.* 1991). Every 33 generations (5 days) we froze glycerol stocks of all 12 populations. Finally, we isolated genomic DNA from the initial and final population samples, and sequenced the whole-population metagenomes to see how the genetic variation changed during the 200 additional generations without recombination. When isolating genomic DNA from these population samples, we took 125 μL of each glycerol stock, then washed and centrifuged the cells twice in DM0 (Davis Minimal medium without glucose) to remove residual glycerol. We then added 100 μL of the washed and resuspended cells into a flask containing 9.9 mL of DM100 (the same medium used in the LTEE, except with a glucose concentration of 100 μg/mL to yield more cells), and we plated the remaining 25 μL onto a tetrazolium arabinose agar plate to make sure that the washing steps did not cause any unexpected population bottleneck.

### Availability of data and analysis scripts

Genome sequence data have been deposited in the NCBI database under the study accession SRP114715, with the respective BioSample accessions SAMN07440982 (donors, recipients, and clones REL4397 and REL4398 from population Ara-3), SAMN07440983 (recombinant clones), and SAMN07440984 (mixed populations sampled at the start and end of the STLE continuation experiment), under BioProject PRJNA396011. All data and analysis code for this project, excluding the genome sequencing read files, are available at the Dryad digital repository [doi: pending publication]. The code for the analysis is also available on GitHub (www.github.com/rohanmaddamsetti/STLE-analysis).

## Results

### Structure of recombinant genomes

Figure 1 summarizes the rich and complex information on genomic changes that occurred during the 1000-generation STLE. Two clones were sampled at random from each of the 12 populations in the recombination treatment, and their genomes were sequenced and analyzed along with those of the donors and LTEE-derived recipients. The odd-numbered clones from each population are shown in Figure 1; the even-numbered clones are shown in Supplementary Figure S2. The genomic sites marked in blue show mutations that distinguish the 12 clones that were used as recipients in the STLE from the ancestor of the LTEE, and which were still present in the genome of a recombinant clone from the STLE. These mutations thus arose during the 7000 generations of the LTEE that preceded the start of the STLE, and they persisted for the 1000 generations of the STLE. The clones from populations Ara+3, Ara+6, and Ara-2 have many more such mutations than the clones from the other nine STLE populations, because the recipient clones used to start these populations came from populations that had evolved hypermutable phenotypes during the early generations of the LTEE (Sniegowski *et al.* 1997; Tenaillon *et al.* 2016). The genomic sites marked in yellow are mutations shared by the K-12 donor strains and the sequenced recombinant clone, but which were not present in the recipient clone used to start that population. These K-12 mutations were thus introduced by intergenomic recombination during the STLE.

**Figure 1.**
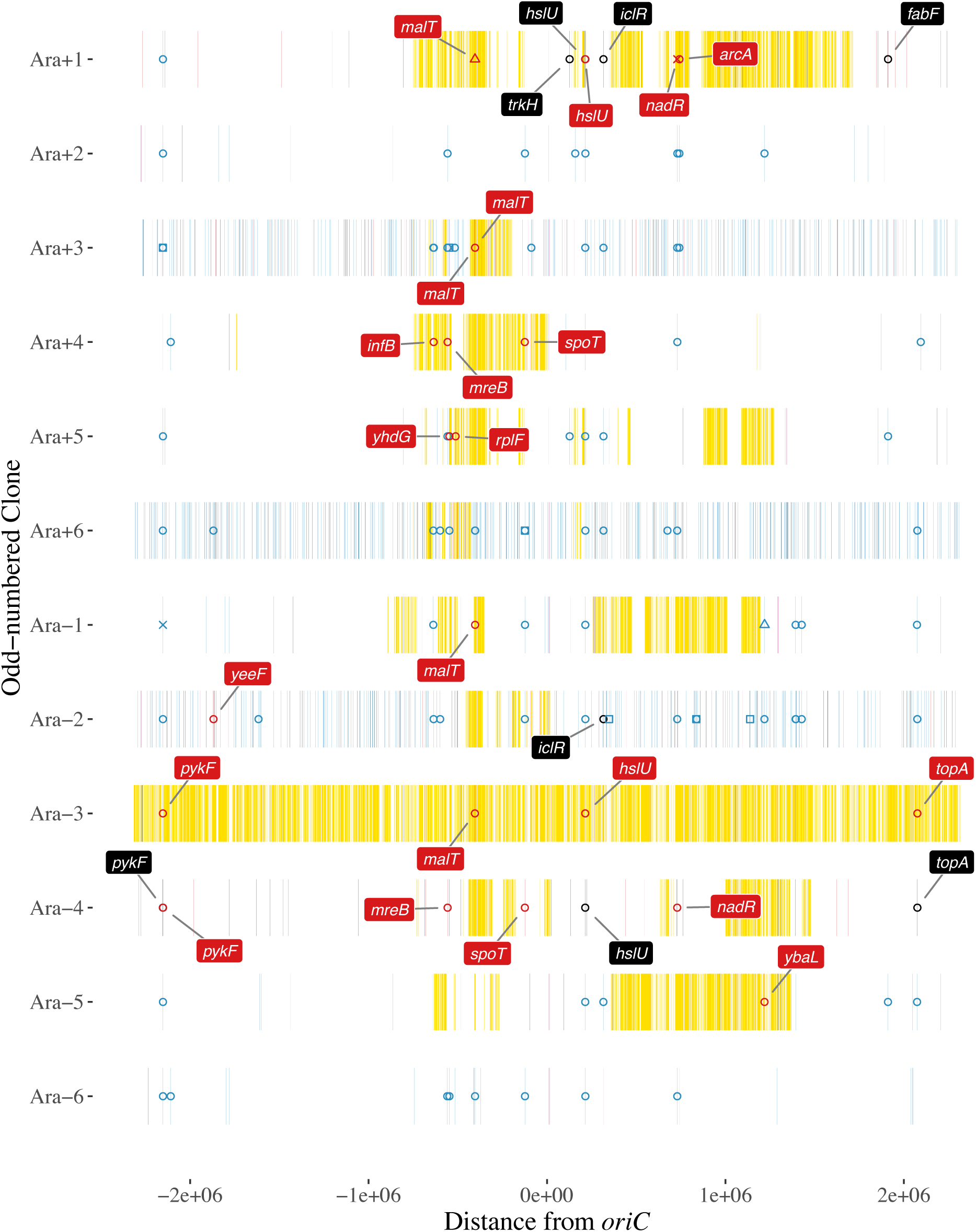
Genome structure of odd-numbered clones from recombinant populations after 1000 generations of the STLE. The REL606 genomic coordinates are shown on the x-axis, centered on the *oriC* origin of replication, and the source populations are shown on the y-axis. Genetic markers specific to K-12 are shown in yellow; mutations specific to the recipient that arose during the LTEE are blue; new mutations that arose during the STLE are black; markers in deleted regions are light purple; and LTEE-derived mutations that were replaced by donor DNA during the STLE are red. The mutations listed in Table 2 that showed strong evidence of positive selection in the LTEE are marked by symbols of the same color. The open circles indicate nonsynonymous point mutations; open squares are synonymous mutations; open triangles are indels; and x-marks are IS-element insertions. Replaced and new mutations in the genes in Table 2 are labeled by their gene names.

**Table 2.**
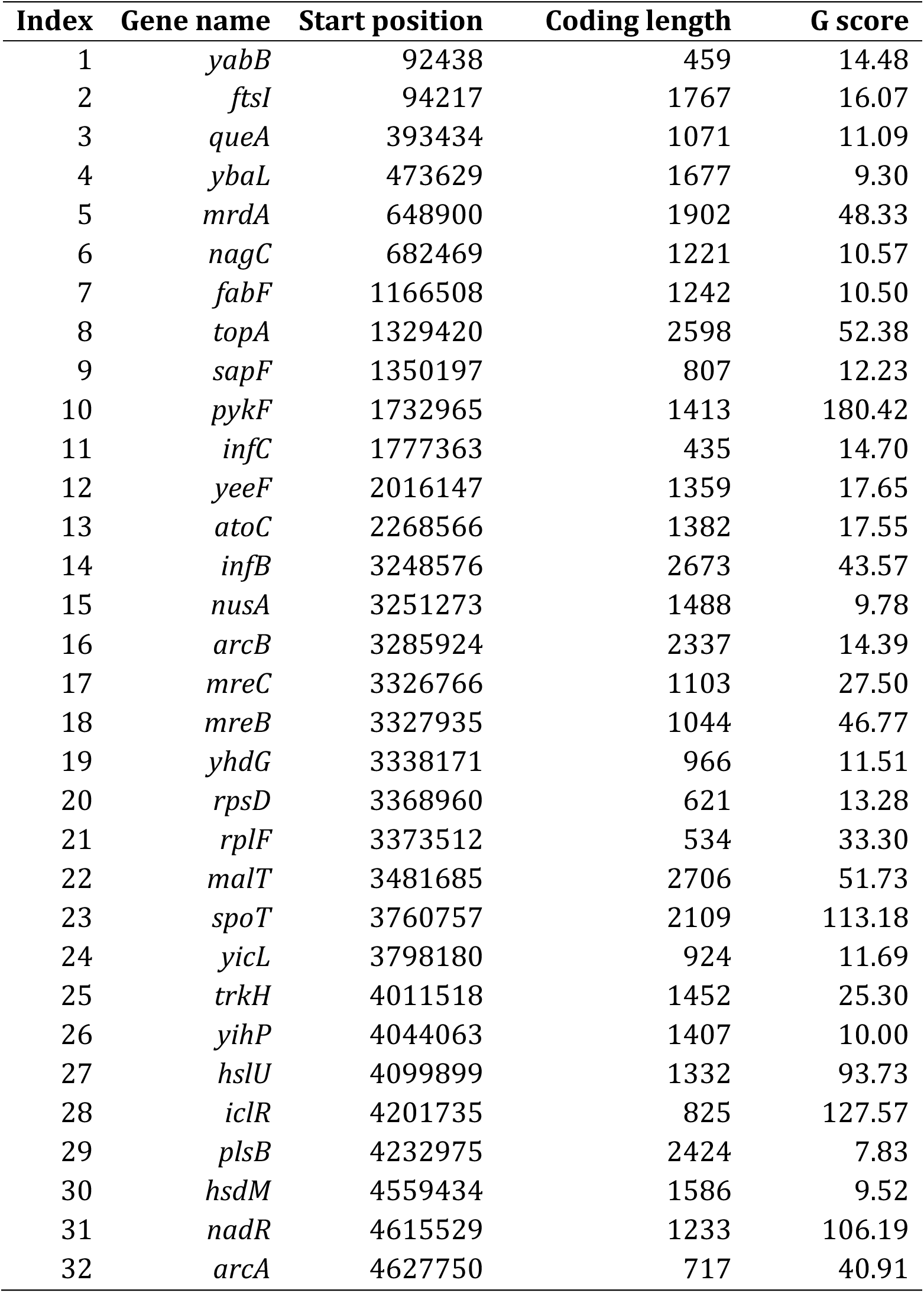
Thirty-two genes under strong positive selection in the LTEE that harbored mutations in at least one of the recipient clones used to start the STLE.

Note: Start position is relative to the reference genome of REL606, and coding length is the number of nucleotides. The G score indicates the strength of evidence for positive selection based on parallel evolution (Tenaillon *et al.* 2016).

These differentially marked sites reveal several features. First, the majority of sites in most genomes derive from the recipients, not from the donor strains. In fact, an individual clone from population Ara-6 (Figure 1) appears to lack any DNA regions that derive from the donors, and both clones from population Ara+2 only have one very short donor segment (~1 kbp in length, barely visible near -1.3 Mbp in Figure 1). Second, there is one striking exception to the above pattern: the genomes of both clones from population Ara-3 are largely comprised of donor-derived DNA (Figure 1). We also sequenced the genomes of two other clones (REL4397 and REL4398) from this same population that were used in a previous study of frequency-dependent selection (Turner *et al.* 1996). They, too, are predominantly K-12, but with many small regions that descend from the LTEE recipient clone (Supplementary Figure S3). Third, in most STLE populations, the pattern of introgression of donor DNA is very similar in the two recombinant clones that we sequenced. However, there are differences in several cases including Ara-6 where, as noted above, one clone appears to lack any donor DNA; Ara+3, where only one clone has any sizable regions of donor DNA; and Ara+4 and Ara-5, where the pairs of recombinant clones share some regions of donor DNA but not others (Figure 1 and Supplementary Figure S2).

Fourth, there is an almost complete absence of donor DNA in all of the recombinant populations (except Ara-3, which has mostly donor DNA) in a span of over 2 Mbp (between about +1.8 to -1.0 Mbp on the circular genomic map). Figure 2A shows this point clearly as the sum of the number of introgressions of donor DNA into the odd-numbered clone from each STLE population, excluding the aberrant Ara-3. This region does not reflect a paucity of mutations that could distinguish the donor-derived and recipient-derived DNA, which are as abundant there as elsewhere in the genome (Supplementary Figure S1). Figure 2A also shows that donor DNA appears to be concentrated in two regions of the recombinant genomes. One region is centered at about -0.5 Mbp on the map, falling off more or less symmetrically on either side; the other donor-rich region peaks at ~1.1 Mbp to ~1.2 Mbp and extends farther to the left.

**Figure 2.**
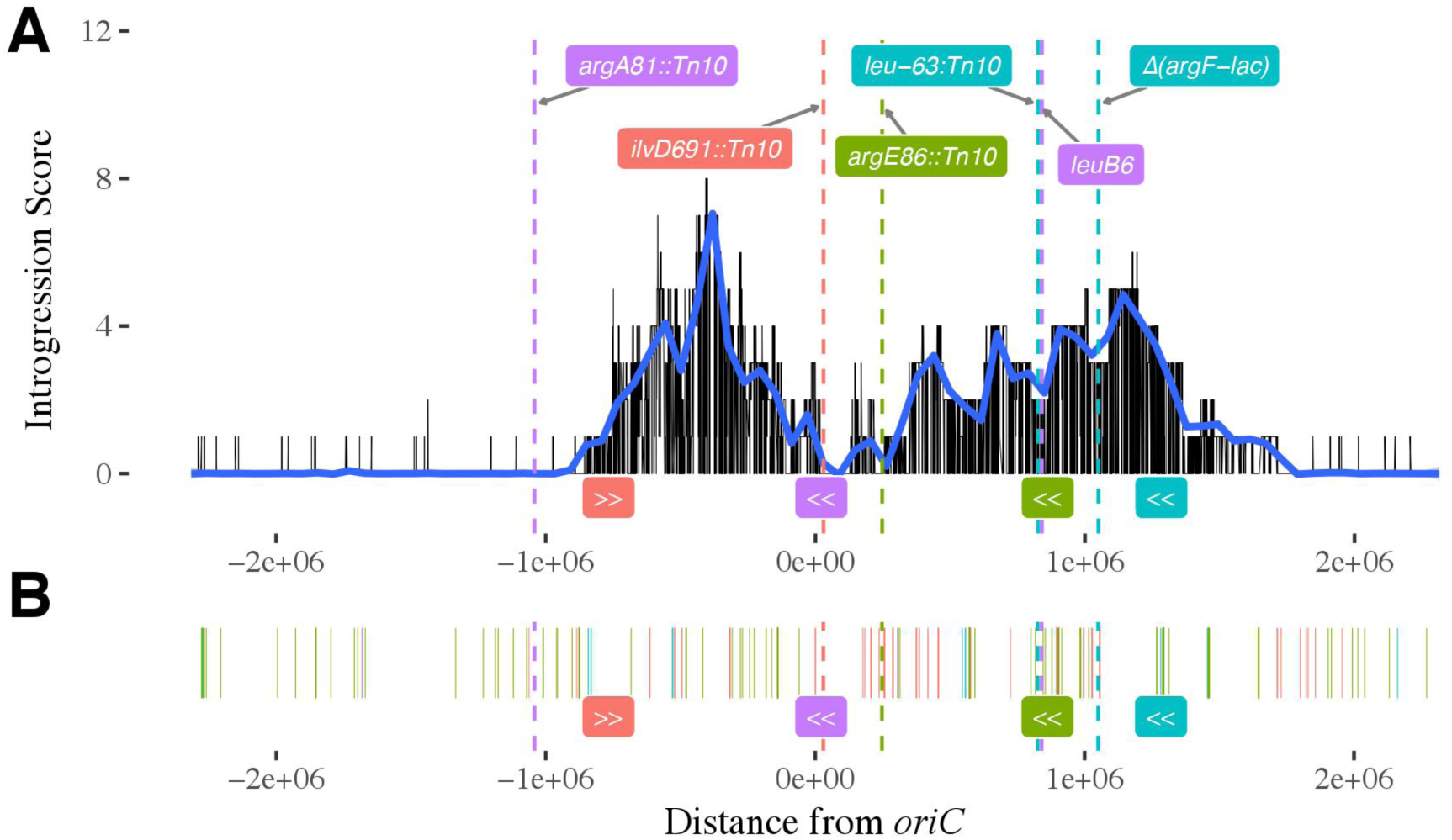
Impact of donors transfer origins and auxotrophic mutations on introgression. (A) The number of parallel introgressions of K-12 genetic markers summed over the odd-numbered STLE clones (omitting the Ara-3 clone, which is almost completely derived from K-12 donor DNA). A natural cubic spline with 100 degrees of freedom, in blue, was fit to these data. The locations of auxotrophic mutations in the donor genomes are shown as dashed vertical lines, and the location and orientation of the *oriT* transfer origin sites are labeled below the x-axis. The markers and transfer origins for REL288 are shown in red, for REL291 in green, for REL296 in teal, and for REL298 in purple. (B) Auxotrophic mutations do not cause a complete barrier to introgression. The location of donor-specific markers found in the odd-numbered recombinant clones (including Ara-3) are labeled with the same colors as in panel (A). Note that almost all of the introgressions near an auxotrophic mutation in one donor strain came from a different donor that did not carry that mutation.

In addition to yellow and blue marks that indicate donor-and LTEE-derived mutations, respectively, Figure 1 also has some black, light-purple, and red marks. Black marks indicate mutations that do not exist in either the donor pool or the recipient clone that was used to start a given population. These marks therefore indicate new mutations that arose during the 1000 generations of the STLE. Not surprisingly, there are many more new mutations in the three populations founded by the hypermutable recipient clones. The light-purple marks indicate regions that were deleted from the recombinant genomes. Red marks indicate mutations present in the LTEE-derived recipient clone but absent from the STLE-derived recombinant clone. These marks imply that the mutations that evolved at those sites during the LTEE were effectively “erased”, being replaced by donor alleles during the STLE. Most of these replaced mutations are surrounded by yellow, showing that they resulted from recombination. However, some of them are not surrounded by yellow; for example, the even-numbered Ara+3 clone has several red marks but no yellow marks (Supplementary Figure S2). Nonetheless, these isolated red marks also probably reflect recombination, not mutation, given the experimental treatment. In particular, if there were no donor-specific markers closely adjacent to a small region of introgressed DNA, then we could not unambiguously assign a segment as donor-derived.

#### Probable beneficial mutations

Figure 1 also has symbols and, in some cases, labels showing the names of certain genes that underwent changes in a recombinant clone. (Supplementary Figure 2 shows the same for the other recombinant clone from each population.) The mutations are marked by symbols that are colored in a similar manner to the lines: blue symbols indicate mutations that were present in the LTEE-derived recipient and retained by the recombinant clone; black symbols are new mutations in the recombinant that were not present in either the recipient or the donor; and red symbols are mutations that were present in the recipient but absent from the recombinant, because they were replaced by donor DNA.

A total of 32 different genes are labeled in one or more recombinant populations. They have been called out because they were probable targets of positive selection under the conditions of the LTEE. Table 2 provides additional information on each of them. These genes were previously identified (along with others that did not undergo any changes in the recombinant clones in our study) by sequencing a total of 264 genomes from the 12 LTEE populations at various times through 50,000 generations, and finding that they had accumulated an unexpectedly large number of independent nonsynonymous mutations in lineages that had not become hypermutable (Tenaillon *et al.* 2016). The *G* scores shown in Table 2 indicate the strength of the evidence for excessive parallelism in a gene, relative to the length of its coding sequence. For some of them, genetic manipulations and competition assays have directly confirmed that mutations indeed improve fitness under the conditions of the LTEE (Barrick *et al.* 2009; Khan *et al.* 2011).

The STLE’s 1000-generation duration is short relative to the 50,000 generations of the LTEE, and so we might not expect to see many new beneficial mutations rising to high frequency in these genes. However, we see many examples including four in population Ara+1 in the *fabF, trkH, hslU*, and *iclR* genes and three in population Ara-4 in the *top A, pykF*, and *hslU* genes (Figure 1). These are also the two non-mutator populations that underwent the most replacements of LTEE-evolved mutations by donor alleles (red hash marks), although we do not know whether this relation is coincidental or meaningful. In the next section, we consider the fate of the presumptively beneficial mutations that were present in the LTEE-derived recipients at the start of the STLE.

#### Possible sources of variation in introgression across genomic regions, and the fate of previously evolved beneficial mutations

What is the source of the variation across the genome in the extent of introgression of the Hfr donors DNA into the recombinant populations? There are several distinct hypotheses that rely either on differences in the propensity for genomic regions to be transferred by the donors or on the fitness effects of integrating different regions into the recipients chromosome. These hypotheses are not mutually exclusive, and so two or more of them may contribute to the observed patterns of introgression (Figures 1 and 2). Hypothesis 1: Some regions of donor DNA were transferred more often than other regions, leading to overrepresentation of the former regions in the recombinant genomes. Hypothesis 2: Some regions of donor DNA contained alleles that were beneficial to the recipient, leading to overrepresentation of those regions in the recombinant genomes. Hypothesis 3: Some regions of donor DNA contained alleles that were deleterious to the recipient, leading to underrepresentation of those regions in the recombinant genomes. This hypothesis can be subdivided into two variant hypotheses. According to Hypothesis 3A, the donor alleles were maladaptive regardless of the beneficial mutations that arose during the LTEE. According to Hypothesis 3B, the donor alleles were maladaptive specifically because the recipient genomes had acquired beneficial mutations in those regions during the 7000 generations of the LTEE that preceded the STLE.

By way of background, it is should be noted that *E. coli* K-12 (the strain that gave rise to the Hfr donors) and B (the strain from which the recipients derive) are fairly closely related, at least as far as *E. coli* strains go. These two source strains were independently isolated from nature many years ago (Daegelen *et al.* 2009). However, about half of their shared genes encode proteins that have identical amino-acid sequences (Studier *et al.* 2009). On the other hand, several hundred genes are present in only one or the other strain, including so-called “genomic islands” that are thought to have been acquired by horizontal gene transfer in the phylogenetic networks leading to one or the other strain (Studier *et al.* 2009). In addition to these more or less ancient differences, the four K-12 donors were deliberately modified by transposon mutagenesis to make them auxotrophic (for different amino acids in the donors) and by introducing the F plasmid (at different locations in the donors) into their chromosomes to make them Hfr (high-frequency recombination) strains; and, as described above, the B-derived recipients accumulated beneficial mutations during the LTEE. Sequence divergence and mismatch repair are known to reduce recombination efficiency (Vulić *et al.* 1999). However, we found that regions containing introgressed K-12 alleles in the STLE-evolved genomes were no more divergent than regions without K-12 alleles (Kruskal-Wallis test, *P* = 0.9091), nor did the extent of sequence divergence between K-12 and the B-derived REL606 correlate with K-12 introgression in the recombinants (Supplementary Figure S4). Therefore, it does not appear as though sequence divergence had much effect on the introgression of K-12 alleles into the STLE populations.

Hypothesis 2 was, in essence, the original motivation for the STLE, with Souza *et al.* (1997) suggesting that intergenomic recombination with the K-12 donors might increase the rate of adaptation (relative to control populations that evolved asexually) by providing an additional source of genetic variation to the LTEE-derived populations. We lack *a priori* information about what sites in the K-12 donor genomes could provide beneficial alleles to the recombinant populations. However, if such sites exist, then we would expect them to be in those regions where the introgression scores are high (Figure 2). On the other hand, Hypothesis 2 seems unlikely, because Souza *etal* (1997) found that fitness gains were not greater in the recombinant populations than in the control populations, which implies that intergenomic recombination did not increase the supply of beneficial alleles.

Figure 2 shows the inferred location and direction of the Hfr origins of transfer of the four K-12 donor strains as well as the inferred location of their auxotrophy mutations, which bear on Hypotheses 1 and 3A, respectively. With respect to Hypothesis 1, Hfr strains transfer their DNA in a unidirectional manner, and the probability that donor genes are transferred to recipients is expected to decline at greater distances from the origin. In the STLE, the cultures in which the donors and recipients were mixed were placed in a non-shaking incubator at 37°C for one hour (Souza *et al.* 1997); that duration would, in principle, allow the transfer of ~60% of the entire chromosome if a conjugative mating began immediately after the strains were mixed. However, not all matings would begin immediately, shaking may disrupt conjugation, and the efficiency of DNA transfer declines with distance from the transfer origin. The peak in the introgression scores between about -1 and 0 Mbp fits very well with the locations and directions of the *oriT* transfer origin sites for two of the Hfr donors: REL288 and REL298 have *oriT* sites near the edges of this peak that point inward from opposite directions. Most of the second, less-defined peak in introgression scores seems to fit moderately well with the other two Hfr donors, REL296 and REL291, whose *oriT* sites are at about 1.3 and 0.8 Mbp, respectively, with the former transferring in the direction of the peak introgression scores and the latter transferring in the same direction toward the broad shoulder between about 0.1 and 1 Mbp. However, the other shoulder of the second, less defined peak—from about 1.2 to 1.5 Mbp—is not explained by the logic of Hfr donor transfer. We also find donor-specific markers in some recombinant clones near the various donor-specific auxotrophic mutations (Figure 2B), although almost all of these nearby introgressions involved a different donor. Nonetheless, the near absence of introgression along the circular chromosome between approximately 1.8 and -1 Mbp—representing over 40% of the genome—fits quite well with the Hfr donor *oriT* sites and directionality. On balance, then, patterns of introgression provide strong, albeit imperfect, support for Hypothesis 1.

We also found compelling evidence of strong purifying selection at the sites of the auxotrophy mutations in the K-12 Hfr donors. Recall that these mutations mean that the cells cannot produce essential amino acids, and therefore the cells cannot grow and persist in the minimal medium used for the STLE. The two donors with transfer properties that can account well for the introgression peak between about -1 and 0 Mbp have auxotrophic mutations located at positions that would sharpen the peak by limiting introgression at each edge. In particular, REL288 has an auxotrophy mutation in the *ilvD* gene that lies just beyond the *oriT* site for REL298; and REL298 has an auxotrophy mutation in the *argA* gene that lies a short distance past the *oriT* site for REL288. The other two Hfr donors, REL291 and REL296, have auxotrophy mutations in the *argE* and *leuB* genes, respectively, that would contribute to the observed decline in introgression scores on the broader shoulder of the less defined peak from about 0 to 1.5 Mbp on the circular map. (REL298 also has a second auxotrophy mutation in *leuB*, but this gene is very far from its *oriT site* and thus probably not relevant to the observed patterns of introgression.) On balance, we also find support for Hypothesis 3A, whereby selection against the effectively lethal auxotrophy mutations in the donor strains reinforces and sharpens the patterns of introgression generated by the mechanics of gene transfer according to Hypothesis 1.

Hypothesis 3B offers a different selection-based explanation for the patterns of introgression. It rests on the idea that selection should also act against donor alleles in those genes where beneficial mutations arose in the LTEE and were present in a given recipient at the start of the STLE. If this hypothesis were correct, then we would expect to see few, if any cases, where these beneficial mutations were removed and replaced by donor DNA. The evidence in support of this hypothesis is ambiguous, at best, because of the considerable variation among the recombinant clones, in terms of both the proportion of their DNA that comes from the donor strains and the extent to which the LTEE-derived beneficial mutations have been retained or replaced. Also, replacements of LTEE-derived beneficial mutations might simply reflect the recent introgression of donor alleles into the recipients (i.e., shortly before the STLE ended)—alleles that would eventually go extinct if conjugation were stopped. In STLE population Ara-1, only 1 of the 9 presumed beneficial mutations present in the recipient was replaced by donor DNA (in both recombinant clones), but ~22% of the recombinant genomes was donor DNA (Figure 1). This pattern is consistent with Hypothesis 3B. By contrast, consider population Ara+1: 4 of the 5 presumed beneficial mutations in the recipient were replaced, but only ~32% of the two recombinant genomes came from the donors (Figure 1). Across the 12 populations, there is a slight tendency for these presumed beneficial mutations to have been replaced by donor alleles more often than the average genomic site, contrary to this hypothesis. On balance, the evidence does not support Hypothesis 3B.

We can also exclude a hypothetical scenario in which beneficial LTEE-derived alleles in the recipients were generally replaced by K-12 alleles that were as or more beneficial. We examined the recombinant clone sequences to determine whether these replacements reverted the gene to its pre-LTEE ancestral state (i.e., the corresponding sequence in REL606, including the case in which that sequence is identical in REL606 and K-12) or, alternatively, introduced a different allele. As usual, we summarize the results for the odd-numbered final clones, but the results do not differ substantively for the even-numbered clones. We examined alignments of 60 proteins (from both non-mutator and hypermutator clones) containing LTEE-evolved alleles in the recipients that were replaced by recombination with the donors. Of those proteins, 11 changed to the K-12 donor state that differs from the pre-LTEE REL606 state; 37 changed to the K-12 donor state that is identical to the pre-LTEE REL606 state; 8 changed back to the pre-LTEE REL606 state that differs from the K-12 donor state; and 4 changed to new states comprising combinations of K-12, REL606, and new mutations.

Figure 3 illustrates the four cases in which new alleles emerged. The *yghJ* gene in population Ara-1 was evidently affected by at least three recombination events: the whole gene is derived from K-12 except for two regions, (spanning residues 803-824 and 1068-1212 in the alignment) that contain REL606 markers. In an *hslU* allele from Ara+1 and a *pykF* allele from Ara-4, recombination events reverted alleles with evolved mutations to their ancestral states, and the new mutations presumably arose later. As seen in the *yghJ* gene from Ara-1, the *nfrA* gene from Ara+3 contains a hybrid allele generated by intragenic recombination (Figure 3). The recombination event in *nfrA* reverted a W289* nonsense mutation to its ancestral state, but left unchanged a C144R substitution. The K-12 markers present at amino acid 364 and beyond were not introduced, and so this recombination event introduced a segment that was at most 364 - 144 = 220 amino acids, or 660 bp, in length. However, it is also possible that this reversion occurred by a point mutation, given the hypermutability of the Ara+3 recipient. (Figure 1). Sequence alignments of all replaced alleles are provided in Supplementary Data File 1.

In general, it is difficult to discern which replaced LTEE alleles in the hypermutator STLE clones (those isolated from populations Ara-2, Ara+3, Ara-6) were beneficial driver mutations, owing to the large number of quasi-neutral passenger mutations that hitchhiked to high frequency in those populations (Tenaillon *et al.* 2016). In contrast, we are confident that LTEE-derived alleles in genes where mutations repeatedly reached high frequency in populations with the ancestral mutation rate were under positive selection (Table 2). We therefore closely examined the alignments of the 30 proteins containing LTEE-evolved mutations in the recipients that were replaced by recombination in the nine odd-numbered non-mutator clones. Recombination with the K-12 donors changed 5 LTEE-evolved alleles into the donor state that differs from the pre-LTEE state; changed 18 LTEE-evolved alleles into the donor state that is identical to the pre-LTEE state (i.e., the same sequence is present in REL606 and K-12); changed 4 LTEE-evolved alleles back to the pre-LTEE state that differs from the donor state; and changed 3 LTEE-derived alleles into new alleles (Table 3). Eleven of the 22 cases where an LTEE-evolved allele went back to its pre-LTEE ancestral state occurred in genes under strong positive selection in the LTEE (Tenaillon *et al.* 2016; Maddamsetti *et al.* 2017), indicating that many beneficial mutations were removed by recombination with the donors. The new alleles that evolved after recombination had effectively reconstructed the ancestral states in *hslU* in Ara-1 and *pykF* in Ara-4 imply that these reversions of beneficial LTEE-derived mutations by recombination were not adaptive.

**Figure 3.**
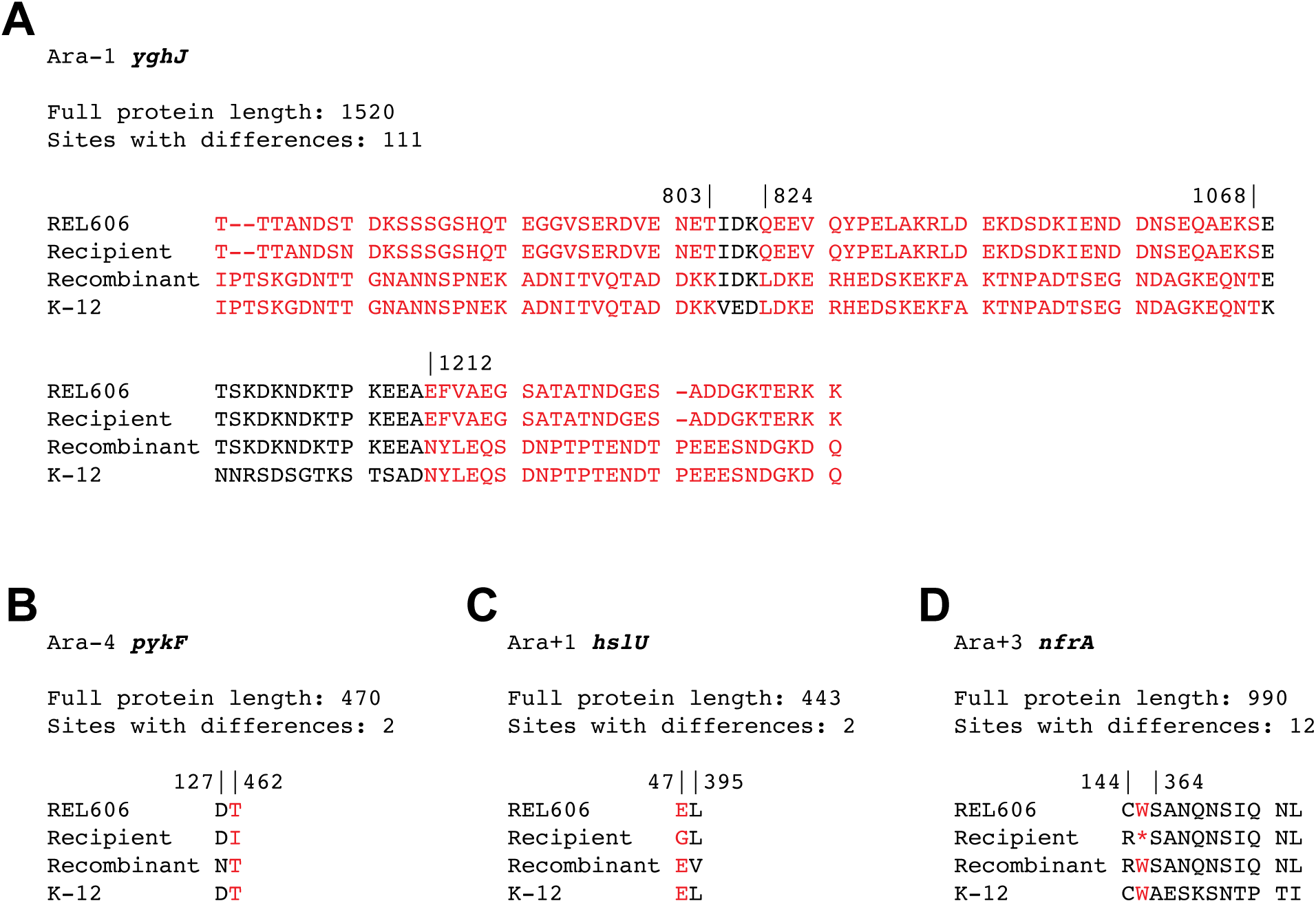
Protein alignments containing new alleles generated via recombination or by subsequent mutation in odd-numbered STLE recombinant clones. Only variable sites are shown. Red columns show K-12 markers introduced by a recombination event. (A) Ara-1 *yghJ*: this locus appears to have experienced at least 3 recombination events. (B) Ara-4 *pykF*: the recombinant is missing a T462I mutation present in the recipient, and it has a new D127N mutation. (C) Ara+1 *hslU*: the recombinant is missing an E47G substitution found in the recipient, and it has a new L395V mutation. (D). Ara+3 *nfrA*: the recombinant lacks the W289* nonsense mutation that is present in the recipient. This putative recombination event did not affect the recipients C144R mutation, nor did it introduce any of the K-12 markers found at residue 364 and beyond. The Ara+3 lineage is hypermutable, and the reversion of the nonsense mutation might have occurred without a recombination event.

**Table 3.** Loci containing nonsynonymous mutations in odd-numbered recipient clones from non-mutator populations that were replaced by donor alleles.

| Population | Gene name | Allelic state of gene after HGT | Strong positive selection in LTEE |
| --- | --- | --- | --- |
| Ara+1 | *arcA* | K-12 | Yes |
| Ara+1 | *hslU* | New allele | Yes |
| Ara+1 | *ycfC* | K-12 (REL606) | No |
| Ara+1 | *yecE* | K-12 (REL606) | No |
| Ara+4 | *infB* | K-12 (REL606) | Yes |
| Ara+4 | *mreB* | K-12 (REL606) | Yes |
| Ara+4 | *spoT* | K-12 | Yes |
| Ara+5 | *rplF* | K-12 (REL606) | Yes |
| Ara+5 | *rpsC* | K-12 (REL606) | No |
| Ara+5 | *yhdG* | K-12 (REL606) | Yes |
| Ara-1 | *malT* | K-12 | Yes |
| Ara-1 | *rpsM* | K-12 (REL606) | No |
| Ara-1 | *yghJ* | New allele | No |
| Ara-3 | *fis* | K-12 (REL606) | No |
| Ara-3 | *hslU* | K-12 (REL606) | Yes |
| Ara-3 | *malT* | K-12 | Yes |
| Ara-3 | *mrdB* | K-12 (REL606) | No |
| Ara-3 | *pykF* | K-12 (REL606) | Yes |
| Ara-3 | *rpsG* | K-12 | No |
| Ara-3 | *tdcR* | K-12 (REL606) | No |
| Ara-3 | *topA* | K-12 (REL606) | Yes |
| Ara-4 | *garP* | K-12 (REL606) | No |
| Ara-4 | *infA* | K-12 (REL606) | No |
| Ara-4 | *mreB* | K-12 (REL606) | Yes |
| Ara-4 | *nadR* | REL606 | Yes |
| Ara-4 | *pykF* | New allele | Yes |
| Ara-4 | *spoT* | REL606 | Yes |
| Ara-4 | *ydfI* | REL606 | No |
| Ara-4 | *ypdC* | REL606 | No |
| Ara-5 | *ybaL* | K-12 (REL606) | Yes |

Note: Allelic state indicates whether the sequence in the recombinant clone is identical to the K-12 but not the REL606 sequence (“K-12”); identical to both the K-12 and the REL606 sequences (“K-12 (REL606)”); identical to the REL606 but not the K-12 sequence (“REL606”); or is a patchwork of K-12, LTEE, and new mutations (“New allele”). See Supplementary File S1 for details.

#### Gene conversion and new mutations

The following analyses focus, for simplicity, on the odd-numbered recombinant clones from the STLE populations that were not hypermutable; however, the even-numbered clones are similar. We noticed that these recombinant clones often had more new mutations than did typical LTEE clones that had evolved for 1000 generations (Tenaillon *et al.* 2016). When we looked for evidence of parallel evolution among these new mutations, we found strong but spurious signals in two genes, *nohB* and *waaQ;* in particular, these genes had multiple identical mutations in multiple lineages. The likely explanation for these parallel changes is gene conversion, in which recombination occurred between non-homologous genes in the K-12 donors and B recipients. To investigate further the possibility of gene conversion, we scored all genes that had three or more new mutations in the same recombinant genome as putative gene conversion events (Table 4). Because so many apparently multi-mutation events occurred, and usually in multiple lineages, we think each case is best explained by a single gene-conversion event, not by multiple mutations in the same gene.

**Table 4.**
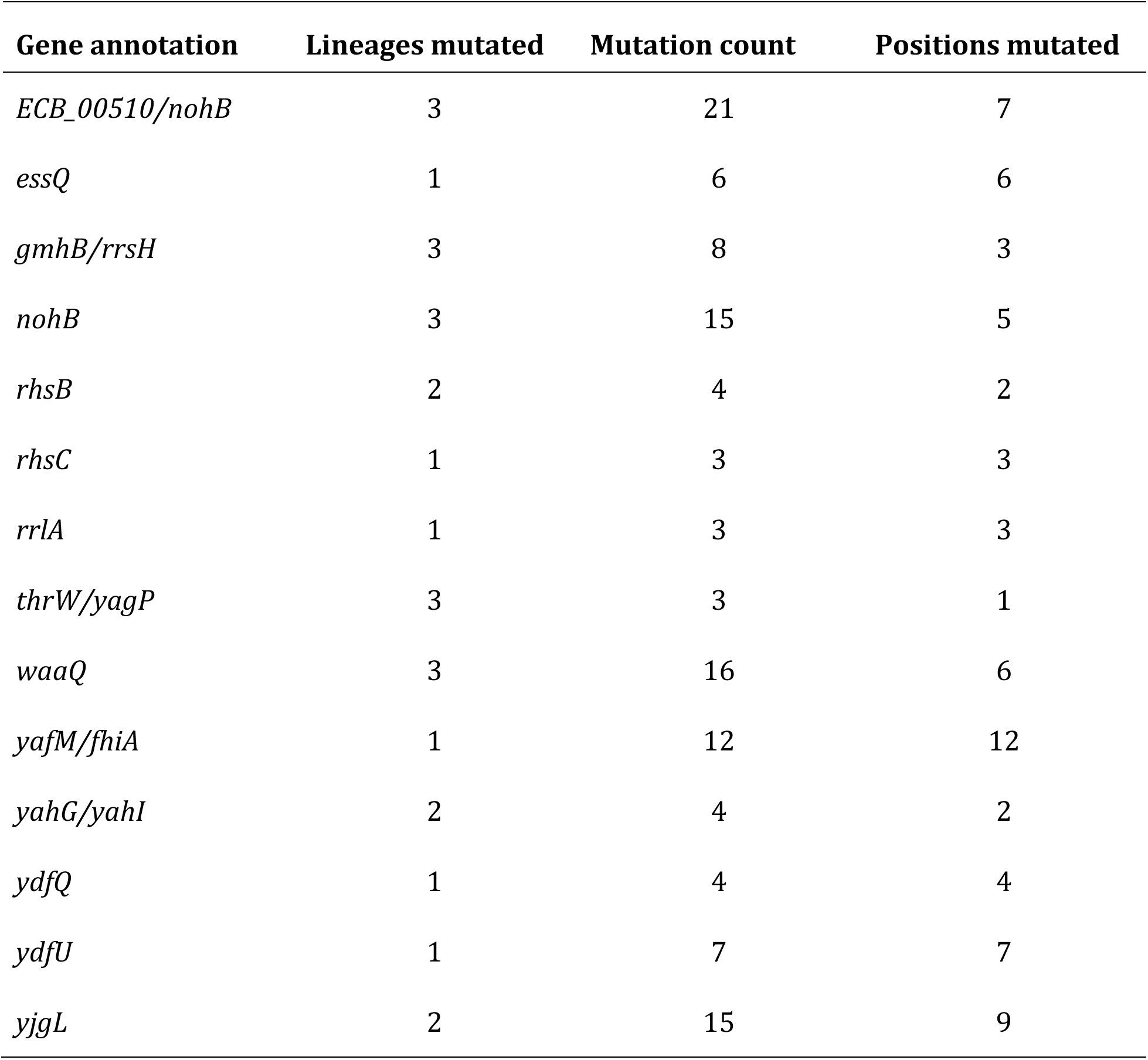
Putative gene-conversion events in recombinant genomes.

| Gene annotation | Lineages mutated | Mutation count | Positions mutated |
| --- | --- | --- | --- |
| *ECB_00510/nohB* | 3 | 21 | 7 |
| *essQ* | 1 | 6 | 6 |
| *gmhB/rrsH* | 3 | 8 | 3 |
| *nohB* | 3 | 15 | 5 |
| *rhsB* | 2 | 4 | 2 |
| *rhsC* | 1 | 3 | 3 |
| *rrlA* | 1 | 3 | 3 |
| *thrW/yagP* | 3 | 3 | 1 |
| *waaQ* | 3 | 16 | 6 |
| *yafM/fhiA* | 1 | 12 | 12 |
| *yahG/yahI* | 2 | 4 | 2 |
| *ydfQ* | 1 | 4 | 4 |
| *ydfU* | 1 | 7 | 7 |
| *yjgL* | 2 | 15 | 9 |

We also used data on positive selection on specific genes in the LTEE (Tenaillon *et al.* 2016) to ask whether new nonsynonymous mutations in the STLE tended to occur in the same genes. Genes affected by nonsynonymous mutations resulting from putative gene conversions had a mean *G*-score of 0, while the genes harboring all other new nonsynonymous mutations had a mean *G*-score of 33.76. This difference, though only marginally significant (two-sided Welchs *t*-test, *p-*value = 0.035), suggests that nonsynonymous mutations that arose during the STLE were under stronger positive selection than those that occurred by non-homologous recombination.

One of our most puzzling findings is that many LTEE-derived mutations, including some that were almost certainly beneficial, were lost in the recombinant clones from the Ara+1 and Ara-4 STLE populations (Figure 1). These genes were among those under strong positive selection for new mutations in the LTEE (Tenaillon *et al.* 2016), and the STLE environment was almost the same as the LTEE. The Ara+1 and Ara-4 STLE populations account for 12 of 30 nonsynonymous replacements, and 14 of 16 new nonsynonymous mutations after excluding the multisite gene-conversion events. This association suggests that the loss of beneficial mutations to recombination in these lineages also led to stronger selection for new beneficial mutations elsewhere in the genome.

In contrast, the Ara-3 STLE population had 8 nonsynonymous replacements, but no new nonsynonymous mutations to compensate. The Ara-3 recombinants are the only ones with genomes that derive primarily from the K-12 donor strains (Figure 1, Figure S2). Also, this population underwent unexpected changes in its ecology, which led to a substantial decline in its fitness relative to the common competitor used in the STLE (Souza *et al.* 1997) and the emergence of a negative frequency-dependent interaction between different recombinant genotypes (Turner *et al.* 1996). These striking changes in genetic background and ecological context might have substantially altered the genetic targets of selection in this population.

#### No characteristic distribution of lengths of recombinant segments

We examined the distributions of recombinant segment lengths to see whether conjugation left a consistent signature in this respect. The left column of Figure 4 shows the distribution of lengths of DNA segments derived from the K-12 donor in the recombinant clones, and the right column shows the length distribution of the B-derived segments (using the odd-numbered clones). To explore the possibility of characteristic segment lengths, we excluded Ara-3 (which was almost entirely donor-derived), Ara-6 and Ara+2 (which had little or no donor DNA), and the three mutator lineages (Ara-2, Ara+3, Ara+6) because DNA repair processes also affect recombination. Even focusing on just the remaining six lineages, the recombinants show significant heterogeneity in the length distribution of their donor-derived segments (Kruskal-Wallis rank sum test, chi-squared = 27.297, df = 5, *p* < 10^−4^). As noted before, K-12 specific markers occur densely over the REL606 reference genome (Supplementary Figure S1), and so the distribution of markers cannot account for this heterogeneity. It is unclear what accounts for this variability across the recombinant lineages. It might reflect differences in the initial recipient genotypes, including their receptivity to conjugation and recombination, or early events in the history of the STLE populations that affected their subsequent receptivity to these processes.

**Figure 4.**
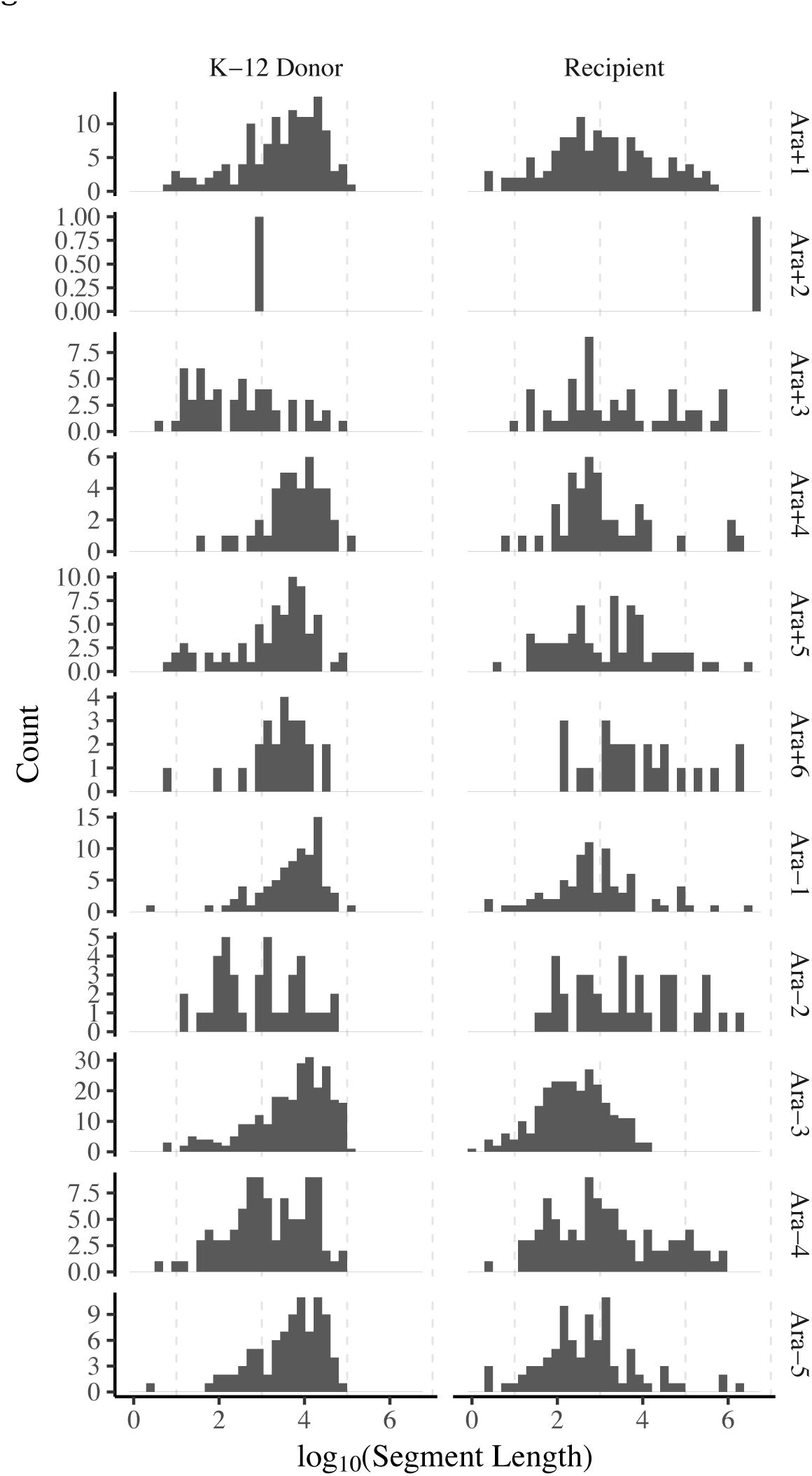
The length distributions, shown on a logarithmic scale, of DNA segments derived from donors (left column) and recipients (right column) in the odd-numbered recombinant genomes.

#### Rate of genomic change due to recombination versus mutation

Table 5 compares the number of synonymous mutations introduced by recombination to those caused by mutation. These data exclude probably spurious inferences of synonymous mutations introduced by gene conversion events (Table 4). We divided new synonymous mutations based on whether they were found in recipient-or donor-derived segments. Even if we exclude the synonymous mutations in the donor-derived segments, the rate at which the non-hypermutable recombinant lineages accumulated synonymous mutations over the 1000 generations of the STLE was several times higher than the corresponding rate during the LTEE (Wielgoss *et al.* 2011; Wielgoss *et al.* 2013; Maddamsetti *et al.* 2015b; Tenaillon *et al.* 2016). This higher rate presumably reflects some mutagenic effect of recombination, such as error-prone repair of double-strand breaks during integration of donor DNA into a recombinant genome. The fact that there are almost as many new synonymous mutations in the donor-derived as in the recipient-derived segments, despite the smaller cumulative target size of the former regions, also supports that interpretation. In any case, the vast majority of synonymous variation that arose during the STLE was introduced through recombination, not by mutation. Twenty of the 24 sequenced recombinant clones acquired hundreds, thousands, and even tens of thousands of synonymous mutations by recombination, whereas no clone, even in the mutator lineages, had as many as 20 synonymous mutations (Table 5). The four exceptional recombinant clones are those that, as described earlier, acquired little or no donor DNA (including both clones from STLE population Ara+2 and single clones from Ara+3 and Ara-6). Excluding those four atypical clones and others from hypermutable populations (Ara+3, Ara+6, and Ara-2), the ratio of synonymous changes introduced by recombination and mutation is generally well over 1000 (Table 5). A recent estimate of the ratio of mutations in recombinant regions relative to spontaneous mutations in natural *E. coli* populations is ~10 (Dixit *et al.* 2015), and so the intergenomic recombination rate in the STLE was clearly much greater than the recombination rate in nature.

**Table 5.**
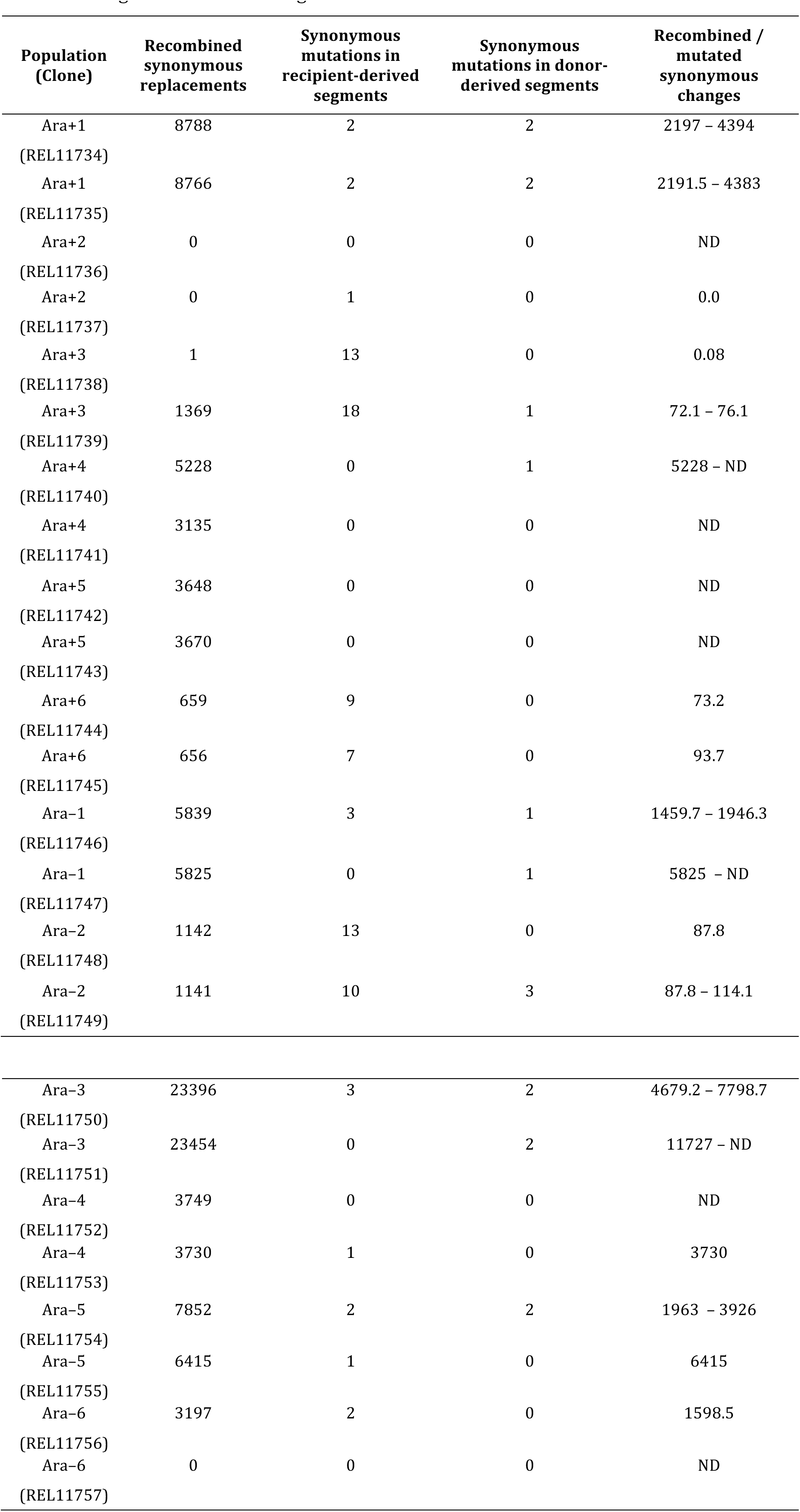
Synonymous changes, including replacements from donors and new mutations, in recombinant genomes after 1000 generations of the STLE.

Note: Ratios of recombined to mutated synonymous changes are calculated with (lower bound) and without (upper bound) synonymous mutations in donor-derived segments. Only one value is shown when there were no synonymous mutations in the donor-derived segments. ND indicates a ratio is not defined because its denominator equals zero.

#### The conjugative Fplasmid persisted in population Ara-3

One unexpected finding is the predominantly donor-derived ancestry of the recombinant clones from STLE population Ara-3 (Figure 1 and Supplementary Figures S2 and S3). We considered the possibility that this ancestry might reflect the reversion of an auxotrophy mutation in one of the K-12 donors, which might have allowed it to persist and perhaps rise to dominance. If that were the case, then we would expect its ancestry to stem from just one of the four donor strains. Instead, the recombinant clones from Ara-3 contain genetic markers from all four donors (as well as the B-derived recipient). In fact, all of the STLE populations (except Ara+2) contain markers derived from at least two of the donors (Figure 5). In any case, the Ara-3 population does not show evidence of being descended from a single donor strain that somehow survived and displaced the recipient population.

**Figure 5.**
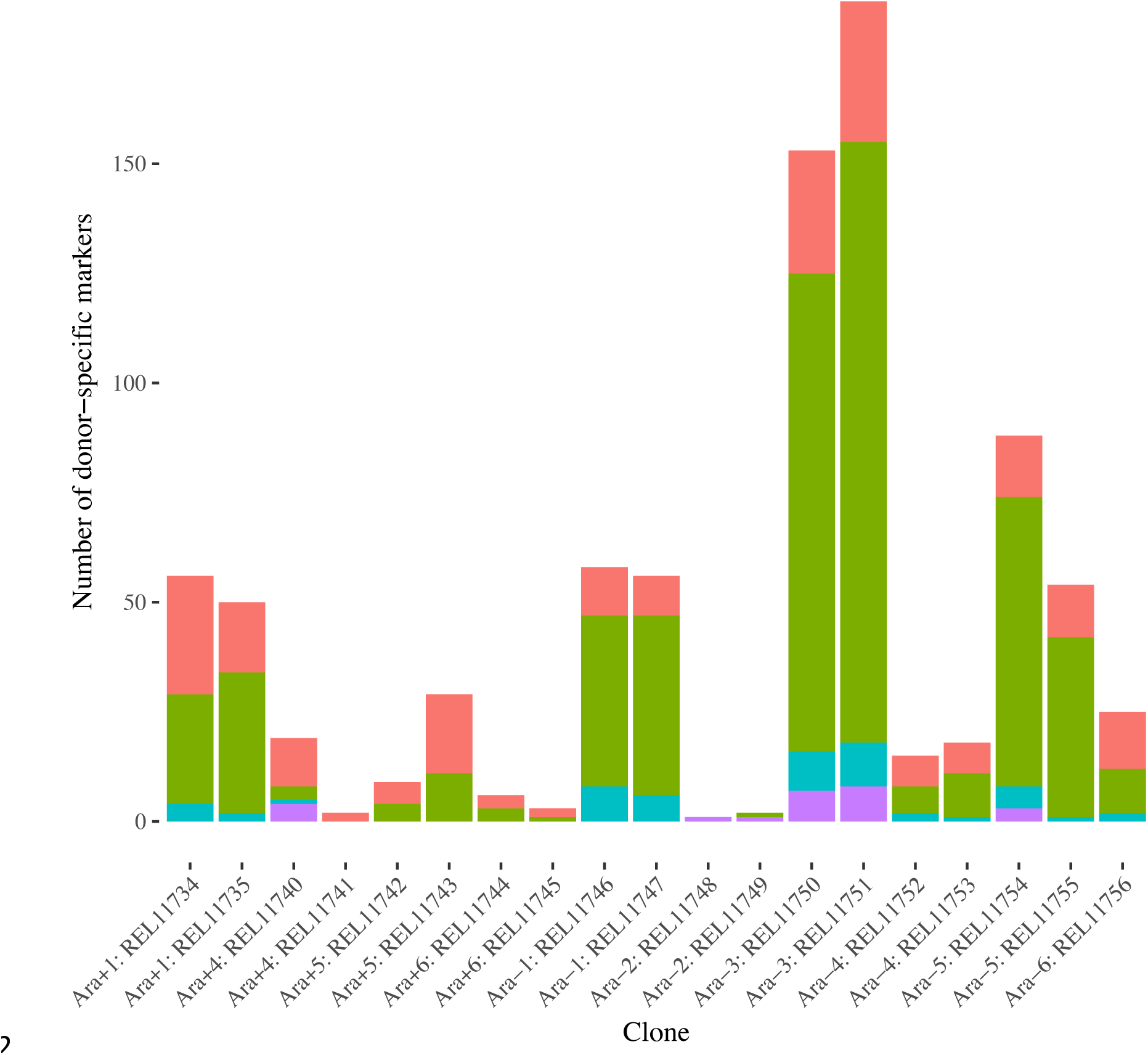
The number and provenance of donor-specific genetic markers found in the recombinant clones. Four clones have very little to no donor DNA, and another has no donor-specific markers.

If the F plasmid persisted in this population, either in a donor-derived lineage or by the conversion of a recipient into a secondary donor, then conjugation might have occurred above and beyond the recombination treatment imposed every fifth day during the STLE. Although no donor-derived lineage appears to have established, even in population Ara-3, it is possible that a recipient was converted to an F plasmid-carrying donor, despite the shaking that occurred after 1 hour of the recombination treatment and was expected to interrupt conjugation (Souza *et al.* 1997). Indeed, more recent studies have found that F-plasmid mediated conjugation readily occurs in *E. coli* at the 120 rpm shaking speed of the STLE, albeit under different conditions (Zhong *et al.* 2010; Wan *et al.* 2011). We found sequencing reads that map to the entire F plasmid in both Ara-3 clones, but not in any other clones we sequenced (Figure 6). We do not know whether these Ara-3 clones contained free copies of the F plasmid or, alternatively, had integrated the F plasmid into the chromosome, which would make them potential Hfr donors. The F-plasmid contains repetitive sequences and insertion elements with homology to multiple locations on the *E. coli* chromosome, making integration a possibility.

**Figure 6.**
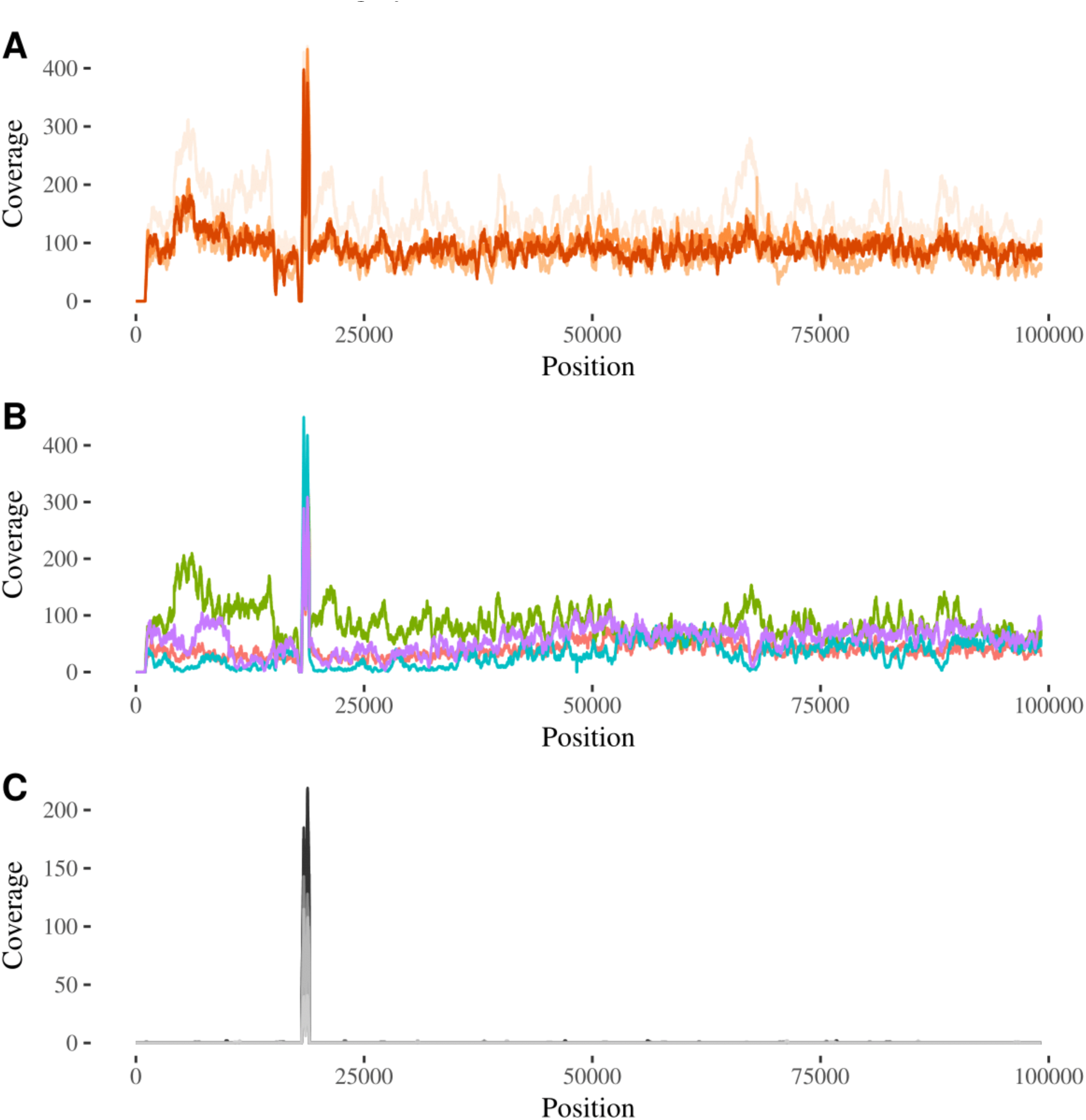
Only Ara-3 recombinant clones contain the F plasmid. Read coverage over the plasmid reference sequence is plotted for each STLE recombinant clone. The spikes in read coverage near the start and near 20 Kbp of the plasmid reference are false-positive reads that map onto repeat sequences in the plasmid. (A) All Ara-3 STLE clones are shown with REL4397 in very light orange, REL4398 in light orange, REL11750 in medium orange, and REL11751 in dark orange. (B) For comparison, the four Hfr donors are shown with REL288 in red, REL291 green, REL296 teal, and REL298 purple. (C) All other STLE recombinant clones are shown in shades of grey.

#### STLE continuation experiment

One potentially confounding factor in our analysis is that some donor segments might have been introduced during one of the final rounds of the conjugation treatment of the STLE, and hence they could be deleterious variants that are present only transiently and destined for extinction. To test that possibility, we restarted the 12 STLE populations from their final samples and then propagated them for an additional 30 days (200 generations) without further addition of the donor strains. This period would thus allow time for recently generated maladapted recombinants to be outcompeted by more-fit members of the respective populations. We then sequenced whole-population samples from the starting and final time points (Table 6). If the introgressed K-12 alleles were deleterious, then we would expect their frequency to decay exponentially over time. Of course, the actual evolutionary dynamics are typically more complicated. For example, the dynamics of large, asexual populations are often driven by the fixation of cohorts of mutations (Lang *et al.* 2013; Maddamsetti *et al.* 2015a; Buskirk *et al.* 2017), and deleterious ‘passenger’ mutations can hitchhike to fixation with beneficial ‘driver’ mutations (Good and Desai 2014). Nonetheless, by measuring how the frequency of K-12 donor alleles changed after stopping conjugation, we can assess whether these alleles were usually evolutionary dead-ends or, instead, persisted by the action of positive selection on introgressed alleles or linked beneficial mutations.

**Table 6.** Population samples used in the STLE continuation and subsequent sequencing.

| Population sample | Population | Generation | Identifying number |
| --- | --- | --- | --- |
| RM3-149-1 | Ara+1 | 1000 | REL4361 |
| RM3-149-2 | Ara+2 | 1000 | REL4362 |
| RM3-149-3 | Ara+3 | 1000 | REL4363 |
| RM3-149-4 | Ara+4 | 1000 | REL4364 |
| RM3-149-5 | Ara+5 | 1000 | REL4365 |
| RM3-149-6 | Ara+6 | 1000 | REL4366 |
| RM3-149-7 | Ara-1 | 1000 | REL4367 |
| RM3-149-8 | Ara-2 | 1000 | REL4368 |
| RM3-149-9 | Ara-3 | 1000 | REL4369 |
| RM3-149-10 | Ara-4 | 1000 | REL4370 |
| RM3-149-11 | Ara-5 | 1000 | REL4371 |
| RM3-149-12 | Ara-6 | 1000 | REL4372 |
| RM3-153-1 | Ara+1 | 1200 | NA |
| RM3-153-2 | Ara+2 | 1200 | NA |
| RM3-153-3 | Ara+3 | 1200 | NA |
| RM3-153-4 | Ara+4 | 1200 | NA |
| RM3-153-5 | Ara+5 | 1200 | NA |
| RM3-153-6 | Ara+6 | 1200 | NA |
| RM3-153-7 | Ara-1 | 1200 | NA |
| RM3-153-8 | Ara-2 | 1200 | NA |
| RM3-153-9 | Ara-3 | 1200 | NA |
| RM3-153-10 | Ara-4 | 1200 | NA |
| RM3-153-11 | Ara-5 | 1200 | NA |
| RM3-153-12 | Ara-6 | 1200 | NA |

Donor-derived alleles increased in frequency in several populations during the STLE continuation (Figure 7), although some populations, most notably Ara+5, showed a mixture of directional changes. Therefore, the introgressed donor-derived alleles do not appear to impose a consistent fitness burden on the recombinant populations. However, we cannot rule out the possibility that some transferred alleles impose a fitness cost, but nevertheless hitchhiked with beneficial mutations elsewhere in the recombinant genomes. Of course, many donor-derived alleles might have increased in frequency as neutral hitchhikers; if a given allele was neutral, then the probability that it would hitchhike to fixation (assuming the STLE continuation was of sufficient duration) would be equal to its initial frequency. Some K-12 donor alleles might even have conferred a selective advantage; the existence of at least a few such alleles is suggested by the tendency for clusters of points to lie above the diagonal.

The continuation experiment and associated metagenomic sequencing also allowed us to infer haplotypes that had fixed before the end of the STLE (Figure 8). Specifically, we identified mutations with frequency equal to 1 at the start and finish of the continuation experiment (corresponding to generations 1000 and 1200, respectively, with respect to the STLE) that were also present in both recombinant clones isolated from a given population. We saw that reversions of highly beneficial LTEE-evolved mutations went to fixation in several of the STLE populations. It is possible that in the three hypermutator populations (Ara+3, Ara+6, and Ara-2), the introgression of donor alleles might have simultaneously removed some deleterious mutations that arose during the LTEE (Wielgoss *et al.* 2013). Nonetheless, it is puzzling to see the recombination-mediated reversion of some clearly beneficial mutations even in non-mutator populations. In some cases (*hslU* in Ara+1, and *pykF* in Ara-4), new mutations became established, presumably after the replacement of the beneficial LTEE-evolved mutations. In other cases *(nadR* in Ara+1 and *spoT* in Ara+4 and Ara-4), beneficial mutations reverted to their pre-LTEE state, and these reversions went to fixation. In general, the locations of parallel introgression events in the inferred LCA of the STLE closely follow the results based on the sequenced clones (Supplementary Figure S5).

**Figure 7.**
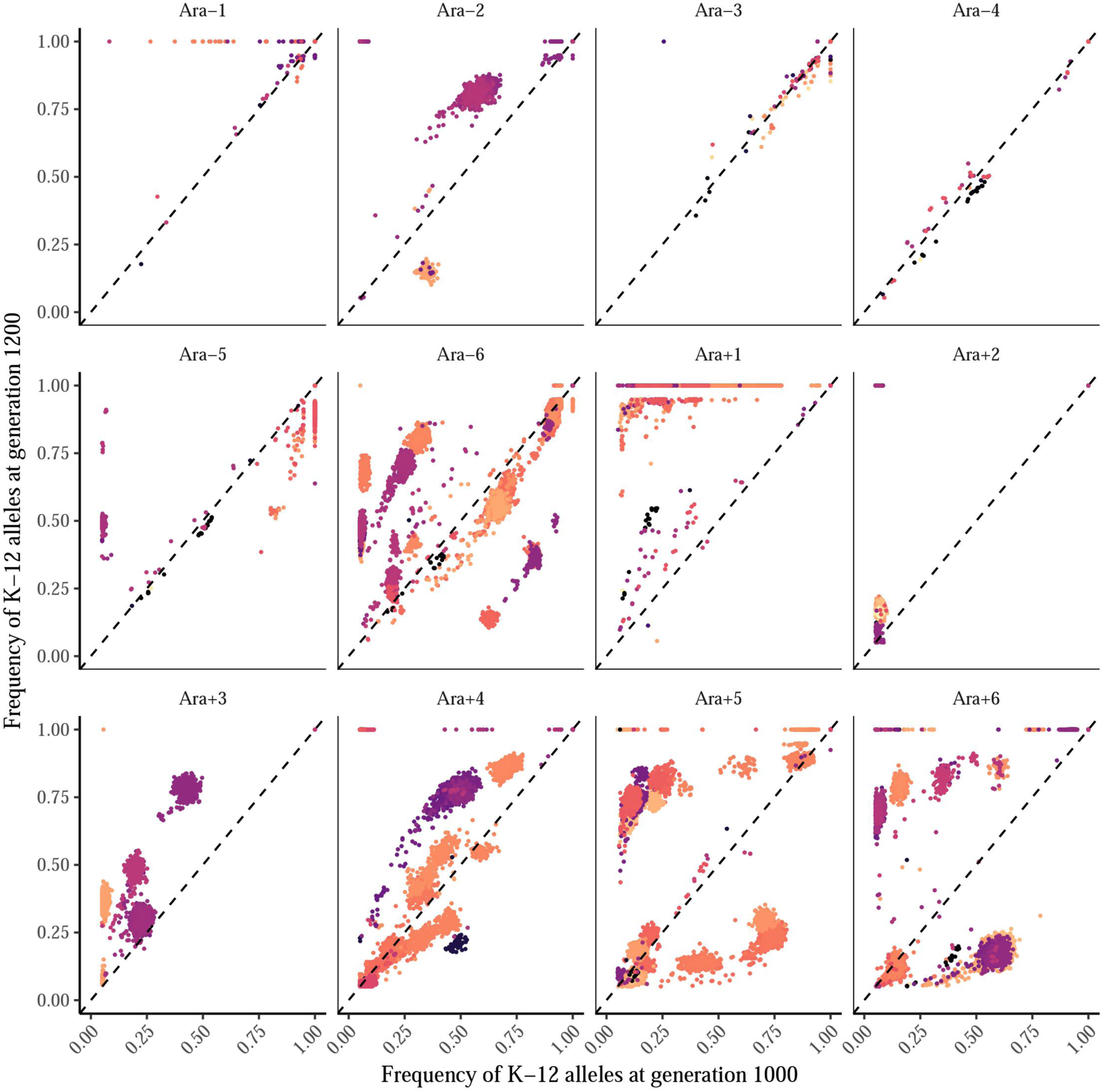
Alleles derived from the K-12 donors tended to increase in frequency during the STLE continuation experiment. Initial and final K-12 allele frequencies are plotted on the x-and y-axis, respectively. Alleles that increased in frequency lie above the dashed diagonal line, and those that decreased in frequency lie below the diagonal. Alleles are colored based on their genomic position, so clusters with the same color probably belong to the same haplotype.

**Figure 8.**
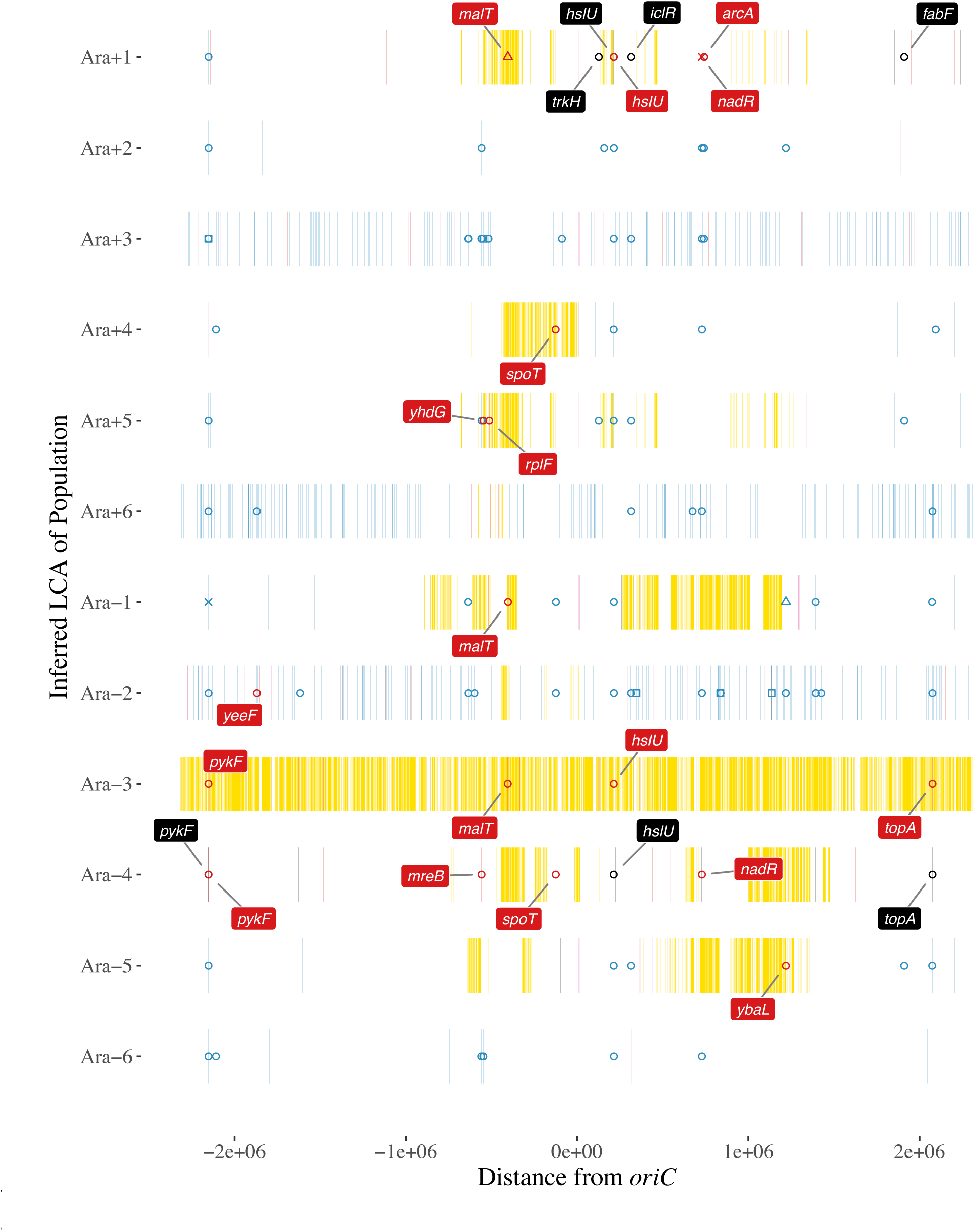
Mutations at 100% frequency in both the initial (1000 generation) and final (1200 generation) samples of the STLE continuation experiment and in both recombinant clones, which we infer to have been present in the last common ancestor (LCA) of each population. REL606 genomic coordinates are plotted on the x-axis, centered on the *oriC* origin of replication, and the 12 replicate populations are shown on the y-axis. K-12 genetic markers are shown in yellow; recipient mutations that arose during the LTEE are blue; new mutations that arose during the STLE are black; markers in deleted regions are light purple; and recipient mutations that were replaced by donor DNA are red. The mutations’ listed in Table 2 with strong evidence of positive selection in the LTEE are marked by symbols of the same color; open circles indicate nonsynonymous mutations, open squares synonymous mutations, open triangles indels, and x marks indicate IS-element insertions. Replaced and new mutations in the genes in Table 2 are labeled by their gene names.

Given that the F plasmid was found in the Ara-3 STLE clones, we looked for its presence in the whole-population metagenome samples corresponding to the start and end of the continuation experiment. The F plasmid was present in the final samples from the continuation experiment in two populations, Ara+1 as well as Ara-3 (Supplementary Figure S6). To get a sense of the frequency of the plasmid in these two populations, and how it changed during the continuation experiment, we compared sequence read coverage of the plasmid and on the chromosome in the initial and final samples. In population Ara-3, the estimated plasmid frequency fell from 75% to 49% during the continuation experiment, whereas in Ara+1 it increased from <1% to 36% (Table 7). Even excluding these two populations, donor-derived alleles tended to increase in frequency in several of the other continuation populations (Figure 7), indicating that plasmid persistence was not necessary for that outcome. Overall, these results indicate that introgressed K-12 alleles did not generally impose a sufficient fitness cost to drive down their frequency, even in the absence of further conjugation.

**Table 7.** Estimated frequency of F plasmids in populations Ara+1 and Ara-3 of the STLE.

| Population | Generation | Average coverage of chromosome | Average coverage of plasmid | Estimated plasmid frequency |
| --- | --- | --- | --- | --- |
| Ara+1 | 1000 | 647.8 | 3 | 0.0046 |
| Ara+1 | 1200 | 686.9 | 250.3 | 0.36 |
| Ara-3 | 1000 | 730.2 | 545.1 | 0.75 |
| Ara-3 | 1200 | 433.1 | 213.6 | 0.49 |

Note: Estimated plasmid frequency assumes one chromosome per cell and one plasmid per plasmid-bearing cell.

Recurrent conjugation over a sufficiently long period means that some neutral, and perhaps deleterious, donor alleles will almost certainly fix in an adaptively evolving recipient population through a ratchet-like hitchhiking process. Suppose that a neutral donor allele is introduced into a genetic background that contains a beneficial mutation undergoing a selective sweep; depending when the donor allele was introduced relative to the sweeping mutation, the donor allele will reach some frequency *p*. Then, the probability that the next beneficial sweep drives that neutral donor allele to fixation is *p*. With probability 1 - *p* the next beneficial mutation occurs on a different genetic background, thereby driving the donor allele to extinction. Nonetheless, the process repeats again and again with each conjugation event. We suggest that this ratchet-like process might explain the some of the unexpected dynamics observed during the STLE.

### Discussion

In this study, we sequenced and analyzed donor, recipient, and recombinant genomes from the Souza-Turner-Lenski experiment (STLE). Two main results are clear. First, we found parallel evolution in the genomic structure of recombinant *E. coli*. This pattern appears to be largely explained by the biases caused by the molecular biology of conjugation, coupled with selection against the effectively lethal auxotrophy mutations carried by the donor strains. Second, there was substantial introgression of donor DNA into most of the recipient populations. Indeed, the rate of recombination was sufficiently high that many beneficial mutations, which had previously evolved during 7,000 generations of asexual evolution in the same environment, were erased by recombination. We estimate that the effective recombination rate, expressed relative to the rate of genomic change by new mutations, was at least 100-fold higher in the STLE than previously estimated in nature for *E. coli* (Dixit *et al.* 2015).

In addition, and to our surprise, we found that one STLE population, designated Ara-3, had predominantly K-12 donor ancestry. The Ara-3 recombinant clones lack all of the LTEE-derived mutations present in their ancestral recipient, but they still have small segments that derive from the *E. coli* B progenitor used to start the LTEE. The effect of recombination was so strong in this population that highly beneficial alleles in the recipient clone at the *pykF, malT, hslU*, and *topA* loci were erased (Figure 8). Moreover, the entire sequence of the F plasmid was present in both of the Ara-3 recombinant clones, whereas it was absent, as expected, from the sequenced recombinant clones from all other populations. (The F plasmid was found at a much lower frequency at the end of the STLE in one other population.) The F plasmid clearly became established in the Ara-3 population, but it is unclear whether it did so as a free plasmid or after being integrated into a recipient chromosome. Several explanations for the F plasmid’s persistence are possible, and they are not mutually exclusive. First, experiments have shown that F+ strains can convert F-cells into F+ cells during a quasi-epidemic of plasmid transmission (Lin *et al.* 2011). If this occurred in Ara-3, then newly infected F+ recipients could transmit the plasmid, along with any donor genes that might hitchhike, to further recipients. Second, a B recipient might have been converted into an Hfr donor and delivered small B segments to a K-12 donor strain that then survived. Third, recombination between the K-12 donor and B recipient genomes might have activated an otherwise latent prophage, leading to virus-mediated transduction in the opposite direction to conjugation. Fourth, a K-12 donor strain might have reverted its auxotrophy mutation, allowing it to grow and persist in the minimal medium of the STLE. It is known that the Tn10-transposon mutagenesis used to construct the donor strains (Wanner 1986) yields unstable genotypes, in which the transposons can move to other locations in the genome. Fifth, one K-12 donor might have recombined with a second K-12 strain (which had perhaps lost its F plasmid and thereby become a recipient) in such a way as to repair the nutritional defect. Sixth, some mutation or mutations in the Ara-3 recipient genome may have allowed for vastly more efficient conjugation and DNA incorporation. However, the two recipient strains with defects in their mismatch repair (Ara+3 and Ara-2) did not have more K-12 ancestry than the other recipients, even though previous research has shown that *E. coli* strains with defective mismatch repair have relaxed DNA homology requirements for molecular recombination (Rayssiguier *et al.* 1989).

Even excluding Ara-3, we observed a great deal of heterogeneity in the amount of introgressed DNA across the STLE populations and in the lengths of donor tracts (Figure 4). Researchers studying natural transformation in other bacterial species have reported that donor segments often cluster into complex mosaic patterns, perhaps generated by long stretches of DNA being disrupted after their uptake or as the result of heteroduplex segregation and correction (Mell *et al.* 2014). Our results accord with these previous reports of fine-scale mosaicism of donor-and recipient-derived regions in recombinant genomes. The lengths of many recombinant segments in the STLE are also consistent with the pervasive transfer of genome fragments ranging from ~40 to ~115 kbp reported in natural populations of *E. coli* (Dixit *et al.* 2015). Both generalized transduction and conjugation can produce such long tracts of introgressed DNA, although the relative contribution of these mechanisms to horizontal gene transfer in nature is unknown. Experiments have shown that the spatial separation of donor and recipient strains during growth on surfaces can suppress conjugation (Freese *et al.* 2014), whereas conjugation can more readily spread genetic material in the well-mixed liquid environment of the STLE.

Evolution experiments with both bacteria and yeast have shown that intergenomic recombination can sometimes speed up the process of adaptation by natural selection (Cooper 2007; McDonald *et al.* 2016). In contrast, the STLE shows that recombination can sometimes act in a manner more analogous to an extremely elevated mutation rate, leading to neutral and even maladaptive changes. Such an effect is not without precedent; for example, plant populations that have evolved resistance to heavy metals found in patchily distributed mine tailings have also evolved selfing to avoid the genetic load of pollen from nearby metal-sensitive populations (Antonovics 2006). The high density of the introduced donors relative to recipients, the high effective rate of recombination, and the fact that the recipients but not the donors had adapted to the STLE environment appear to have created a similar situation, in which non-adapted donor genes “rained down” on the locally adapted, LTEE-derived recipients. The most striking evidence that recombination could sometimes have maladaptive effects in the STLE was the finding that many beneficial mutations were effectively “erased” by replacement with donor alleles that were the same as the LTEE ancestral state, especially in populations Ara+1, Ara-3, and Ara-4. If most donor alleles were neutral or maladaptive in the environment of the STLE, then it is not surprising that the recombination treatment did not speed up adaptation (Souza *et al.* 1997). What is surprising, though, is the extent to which those alleles could evidently invade and replace better-adapted recipient alleles. On the other hand, the fact that most STLE populations did not decline in fitness, despite having some beneficial mutations erased, leaves open the possibility that some other donor-derived segments harbored beneficial alleles that offset the removal of LTEE-derived beneficial mutations. Also, by allowing the STLE populations to evolve for an additional 200 generations without the conjugation treatment, we showed that haplotypes containing donor-derived segments often increased in frequency (Figure 7), contradicting the hypothesis that they could only persist by ongoing recombination.

In any case, it appears that donor genes physically linked to their *oriT* transfer sites often replaced homologous genes in the recipient populations by virtue of the transmission advantage produced by conjugation. This outcome lends credence to the hypothesis that the evolutionary origin of sex might lie in a transmission advantage for recombining genes (Hickey and Rose 1986, rather than an improvement in the efficiency of natural selection. Although our analyses examined the effects of bacterial conjugation, our findings might shed indirect light on the origin of meiosis in eukaryotes. In the STLE, it seems that recombination acted, to a large extent, as a gene drive that gave a transmission advantage to donor DNA (including hitchhikers and, occasionally, the conjugative driver itself, despite efforts to prevent the latter in the STLE). Perhaps, then, eukaryotic sex also originated as a gene drive. If so, major questions remain unanswered including: What evolutionary forces favored the transition to cooperation in the evolution of meiosis, since the products of meiosis are usually fairly divided into gametes? And how did this transition occur, given the existence of selfish meiotic drivers in so many eukaryotic species (Bomblies 2014)?

## Acknowledgments

We thank Neerja Hajela for technical assistance, Valeria Souza and Paul Turner for their prior work on the STLE, and Jeff Barrick for advice on *breseq* and the design of this project. We thank Rosemary Redfield, Joshua Mell, Chris Marx, Valeria Souza, and Paul Turner for valuable discussions at the 2015 Microbial Population Biology Gordon Research Conference. R.M. thanks Debora Marks and Chris Sander for support while writing this paper. This work was supported by a grant from the National Science Foundation (DEB-1451740 to R.E.L.), the BEACON Center for the Study of Evolution in Action (NSF Cooperative Agreement DBI-0939454), a National Defense Science and Engineering Graduate Fellowship (to R.M.), and a dissertation completion fellowship from the MSU Graduate School (to R.M.).

## Supplemental Methods

*Calculation of lengths of donor and recipient segments in recombinant genomes*

Each genome is a list of labeled mutations. First, we initialize a list of “0” with the length of the genome. We keep track of two state variables: the index of the last breakpoint (transition-state) and a Boolean state variable called *in.K12.chunk* that is initialized to FALSE under the assumption that the first genomic segment comes from the B recipient. For every labeled mutation in the genome, we check whether the current mutation has a label that changes the state of *in.K12.chunk*. If *in.K12.chunk* is FALSE and its state changes, then the current mutation is labeled “1-2”. If *in.K12.chunk* is TRUE and its state changes, the current mutation is labeled “2-1”. At the end of the loop, we check our initial assumption that *in.K12.chunk* was FALSE. Because we stored the position of the last transition-state, we check whether the last transition-state in the genome is “1-2”, in which case the first “1-2” transition should be set to “0” because the *E. coli* genome is circular. All sites marked “0” are removed from the genome. We then calculate the differences between the N-1 pairs of transition-state mutations: “1-2” on the left and “2-1” on the right gives the length of a K-12 segment, whereas “2-1” on the left and “1-2” on the right gives the length of a B segment. The two final transition-state mutations are the last and first elements of the list. In this way, we calculate the lengths of segments in a recombinant genome that were derived from the donor and recipient, adjusting for any deletions or insertions that may have occurred in those segments.

**Figure S1.**
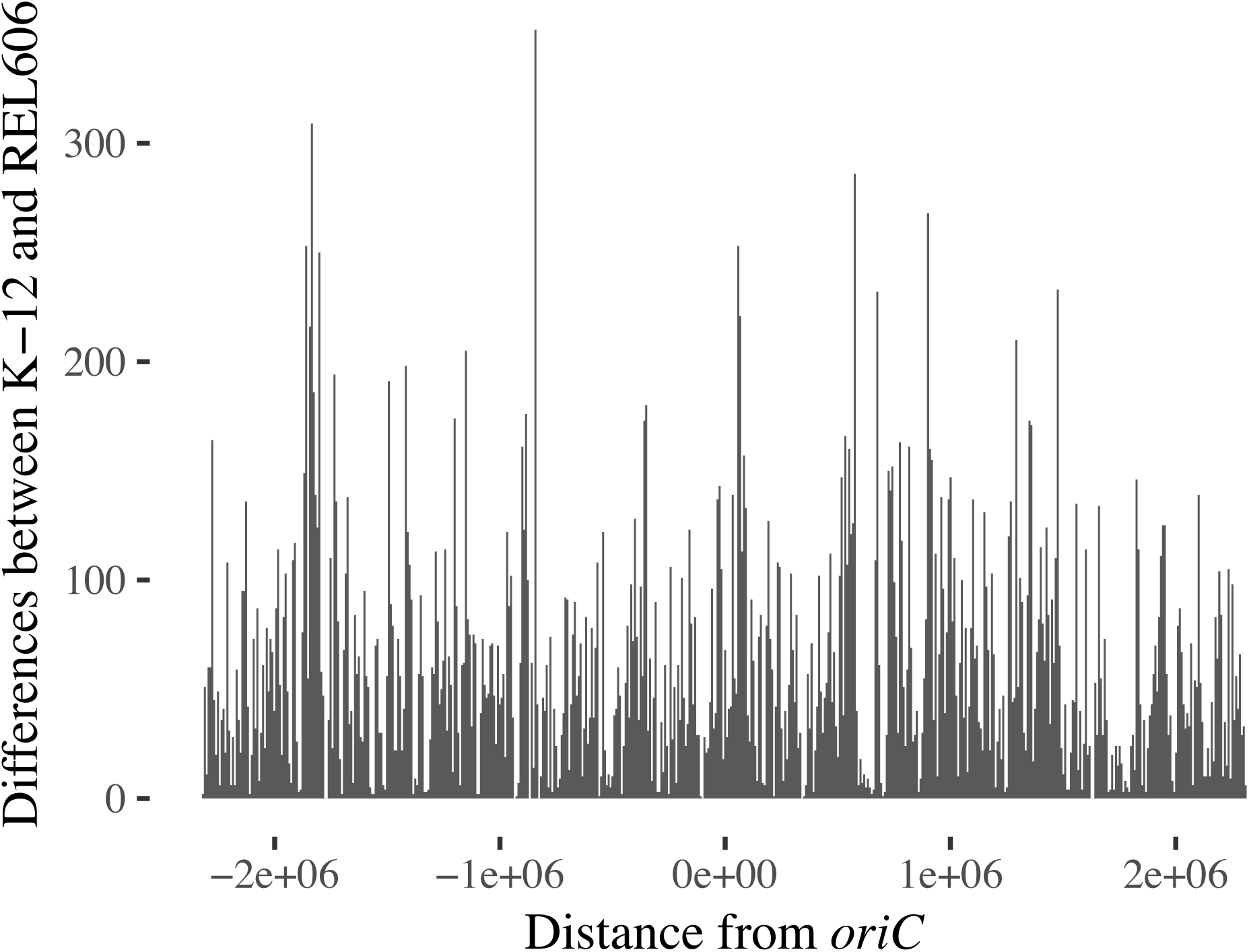
Histogram of the density of K-12 specific marker mutations plotted along the genome coordinates of REL606, the ancestral strain of the LTEE. The numbers were binned over 556 DNA segments that are each 8327 bp in length.

**Figure S2.**
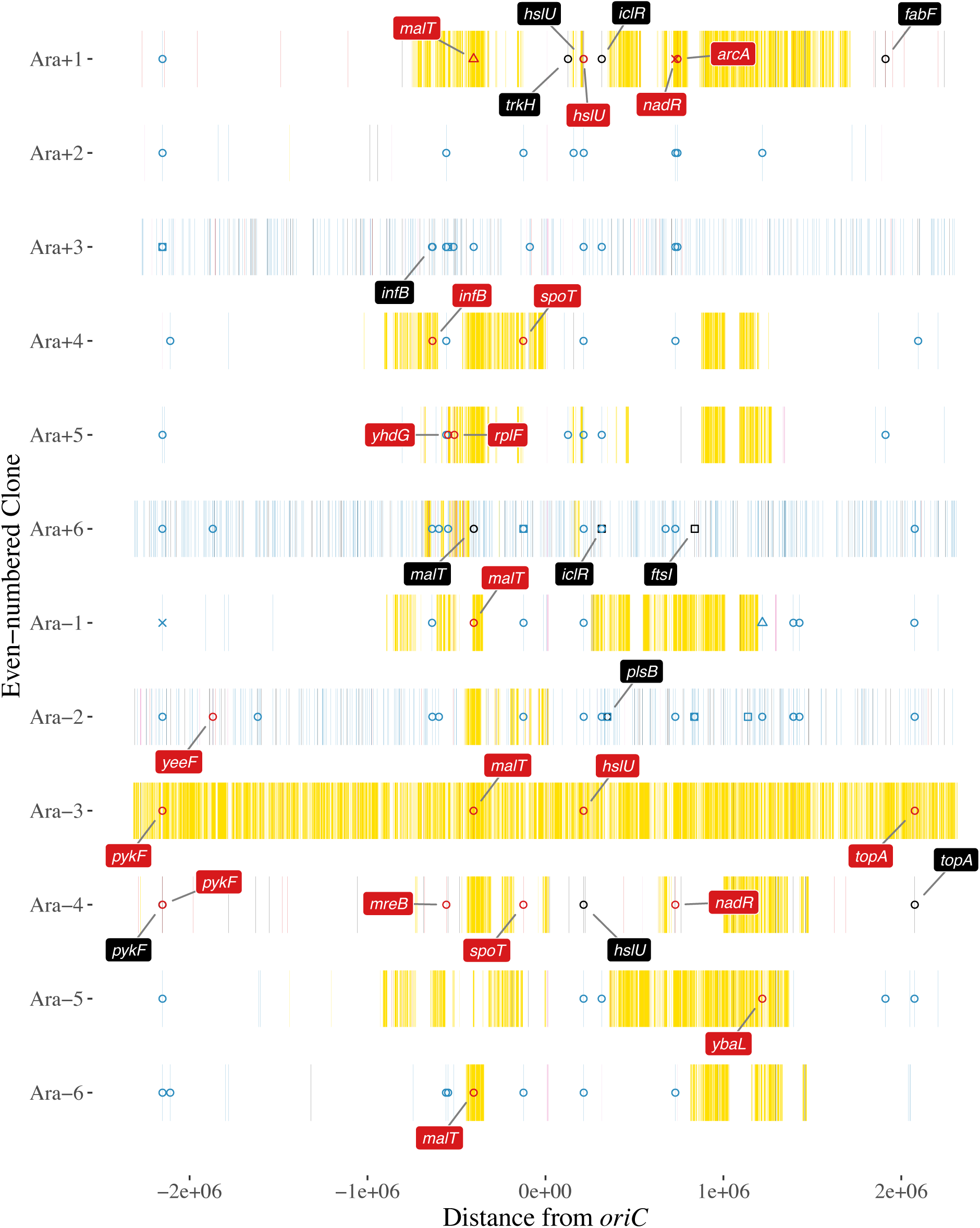
Genome structure of even-numbered clones from recombinant populations after 1000 generations of the STLE. The REL606 genomic coordinates are shown on the x-axis, centered on the *oriC* origin of replication, and the source populations are shown on the y-axis. Genetic markers specific to K-12 are shown in yellow; mutations specific to the recipient that arose during the LTEE are blue; new mutations that arose during the STLE are black; markers in deleted regions are light purple; and LTEE-derived mutations that were replaced by donor DNA during the STLE are red. The mutations listed in Table 2 that showed strong evidence of positive selection in the LTEE are marked by symbols of the same color. The open circles indicate nonsynonymous point mutations; open squares are synonymous mutations; open triangles are indels; and x-marks are IS-element insertions. Replaced and new mutations in the genes in Table 2 are labeled by their gene names.

**Figure S3.**
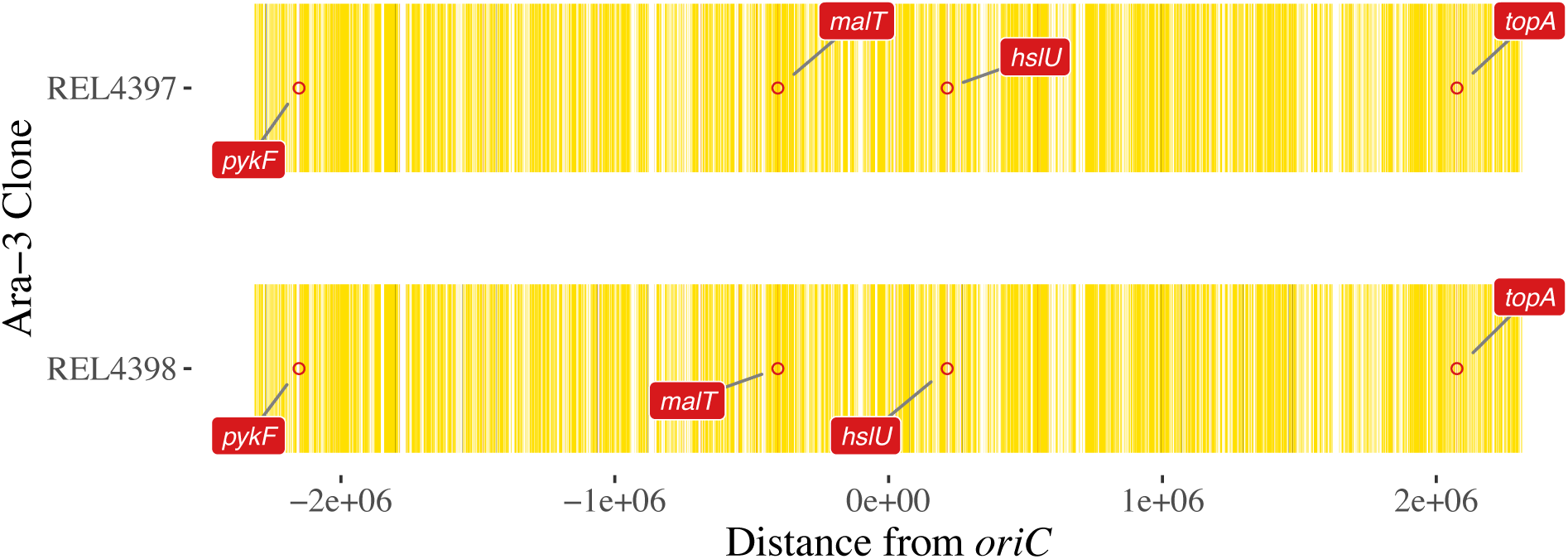
REL4397 and REL4398 are recombinant clones from STLE population Ara-3 that were previously studied by Turner *et al.* (1996). They discovered a cross-feeding interaction between these clones, which puzzlingly had declined in fitness relative to their progenitor during the STLE. The REL606 genomic coordinates are shown on the x-axis, centered on the *oriC* origin of replication, and the source populations are shown on the y-axis. Genetic markers specific to K-12 are shown in yellow; mutations specific to the recipient that arose during the LTEE are light blue; new mutations that arose during the STLE are black; markers in deleted regions are light purple; and LTEE-derived mutations that were replaced by donor DNA during the STLE are red. The mutations listed in Table 2 that showed strong evidence of positive selection in the LTEE are marked by symbols of the same color. The open circles indicate nonsynonymous point mutations. Replaced and new mutations in the genes in Table 2 are labeled by their gene names.

**Figure S4.**
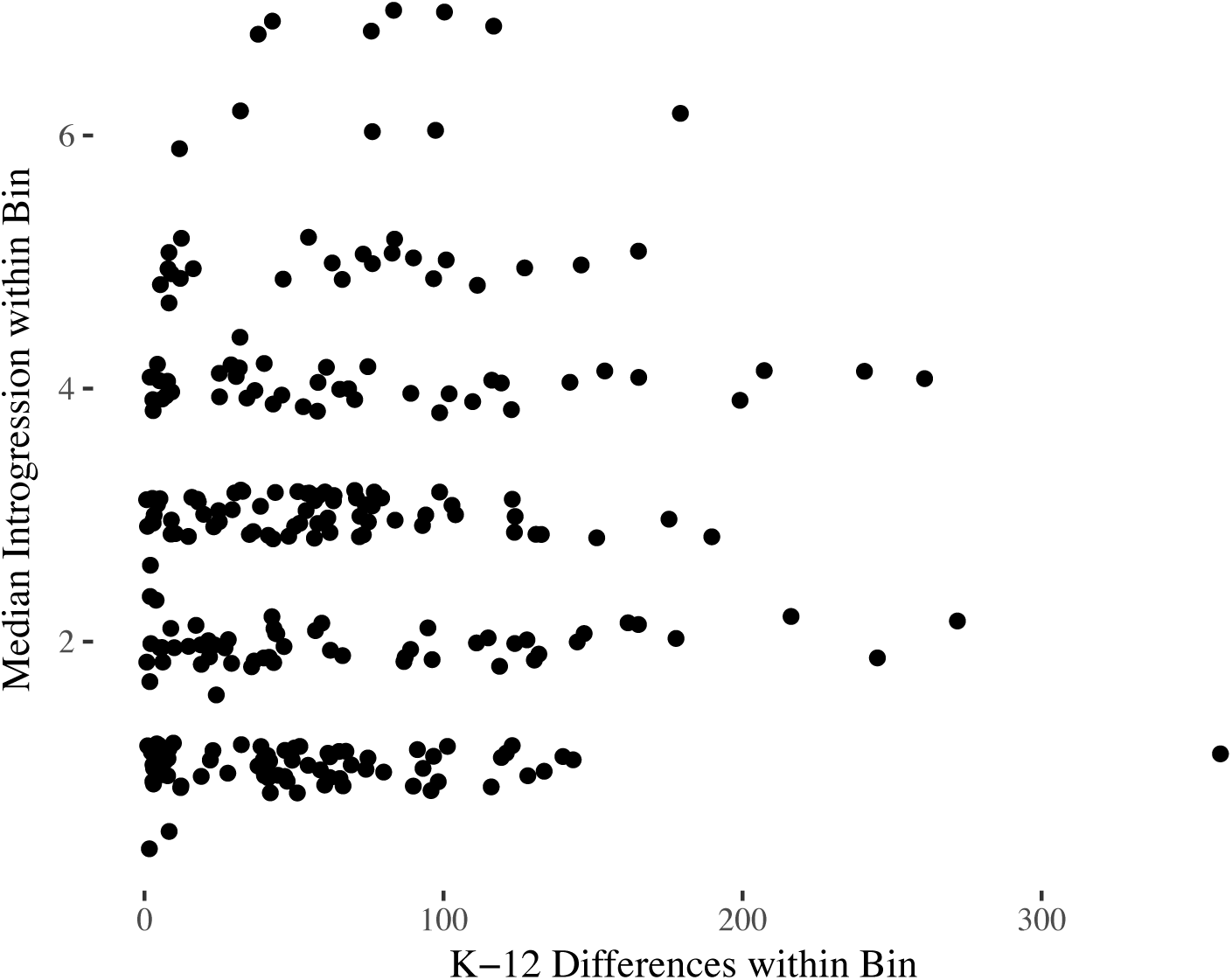
Sequence divergence between donors and recipients does not obviously affect introgression efficiency. The recipients are all recently derived from and closely related to REL606, itself a derivative of *E. coli* B. The REL606 genome is 4629812 bp, which we divided into 556 bins that are each 8327 bp long. For each point, the x-coordinate shows the number of mutational differences between K-12 and REL606 in each segment, and the y-coordinate shows the median number of parallel introgression events (as shown in Fig. 3) within that segment.

**Figure S5.**
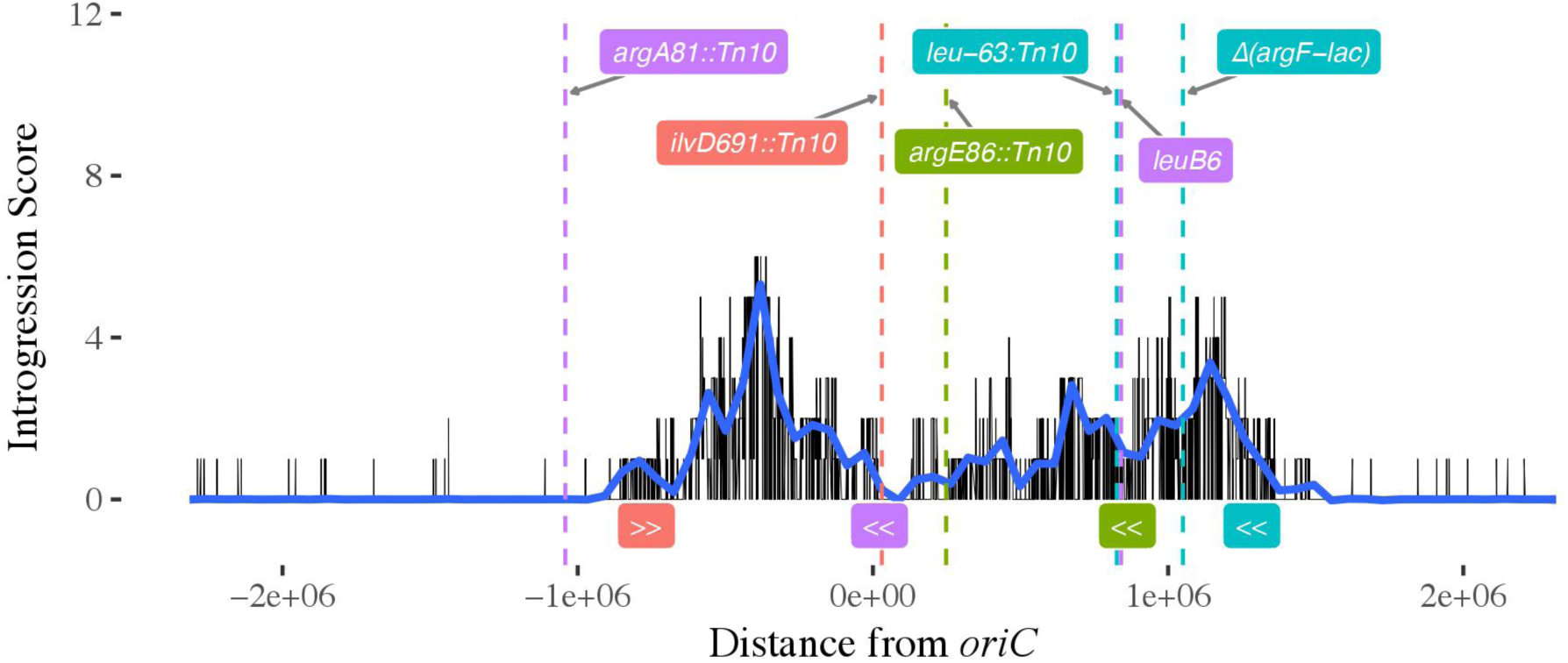
Parallel fixations of K-12 genetic markers during the STLE. K-12 markers found in both recombinant clones and at 100% frequency in both the initial (1000 generation) and final (1200 generation) samples of the STLE continuation experiment were summed over each population (omitting the Ara-3 population which is almost completely derived from K-12 donor DNA). The locations of auxotroph mutations in the donor genomes are shown as dashed vertical lines, and the location and orientation of the Hfr *oriT* transfer origin sites are labeled below the x-axis. The four donors are colored as in Figure 2.

**Figure S6.**
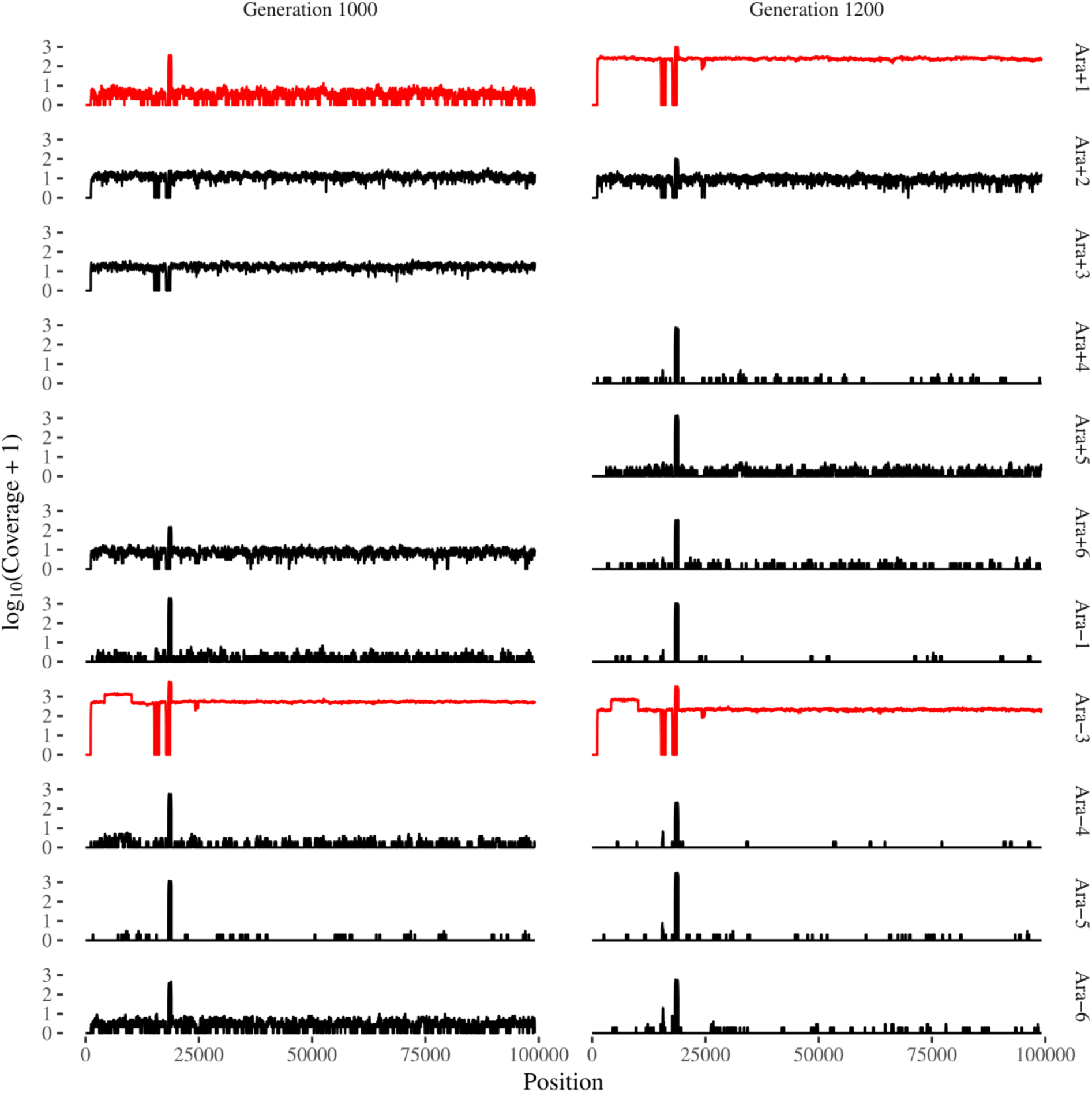
The Ara+1 population at generation 1200 of the STLE continuation contains the F plasmid, as does the Ara-3 population. The Ara+1 and Ara-3 populations are shown in red, and all other populations in black. This plot shows the coverage distribution over the plasmid reference sequence for the initial (generation 1000) and final (generation 1200) samples from the continuation populations. The three blank panels are for samples where *breseq* (version 0.31) indicated insufficient coverage across the F plasmid.

**Supplementary Data File 1**. Whole-protein alignments of proteins encoded by genes in which mutations present in the recipient were replaced by donor DNA in the odd-numbered STLE recombinant clones.

